# Activity of brain stem glucagon neurons are modulated by energy state and encode sex and frequency-dependent negative valence and anxiety

**DOI:** 10.1101/2023.05.11.540425

**Authors:** CB Lopez, M Duran, SA Virkus, E Yadav, K McMichen, J Singh, V Ramsey, S Stocking, KM Habegger, JA Hardaway

## Abstract

The glucagon-like peptide 1 (GLP-1) system has emerged as an important drug target for the treatment of obesity and diabetes. Preclinical and clinical studies demonstrate that the activation of GLP-1 receptors (GLP-1Rs) directly in the brain through overexpression of GLP-1 or GLP-1R agonists produces potent anorexigenic effects, yet the behavioral role and modulation of the *endogenous* GLP-1 producing system in the brain by energy status is unclear. In this study, we examined the anatomical, physiological, and behavioral properties of preproglucagon-expressing neurons in the nucleus of the solitary tract, *Gcg*^NTS^ neurons, which serve as the primary source of GLP-1 in the brain. Using transgenic laboratory mice, we observed no sex differences in the density and distribution of *Gcg*^NTS^ neurons in male and female mice. Fos immunolabeling experiments show that *Gcg*^NTS^ neurons are not significantly activated after intermittent access to palatable food, but the magnitude of Fos activation was linearly related to the amount of food intake in mice provided with *ad libitum* intermittent access to palatable food. Electrophysiological examination of *Gcg*^NTS^ neurons revealed that these neurons show energy-status and sex-dependent changes in neuronal firing and intrinsic excitability. Twenty-four hour food deprivation produced a significant reduction in excitability and firing in male, but not female mice. We then used optogenetics to investigate the causal behavioral role of *Gcg*^NTS^ neurons. High frequency optogenetic activation of *Gcg*^NTS^ neurons using the red light-gated opsin ChrimsonR produced female-specific anxiety-like behavior and real-time place aversion. For feeding, we observed that reversible optogenetic stimulation at high frequencies produced a significant reduction in homeostatic refeeding that did not differ by sex. Using operant conditioning, we found that reversible optogenetic activation of *Gcg*^NTS^ neurons at 20 Hz, but not 5, also reduces appetitive behavior. These data demonstrate that *Gcg^NTS^* neurons exert control over motivation and food-seeking behavior in addition to consumption.

## INTRODUCTION

Impacting more than a third of US adults, obesity is a leading risk factor for cardiovascular disease, hypertension, psychiatric disorders, cancer, and type II diabetes (T2D).^1^ Together, these diseases represent the primary cause of mortality in the US and impose a major economic burden of over $200 billion per year.^2^ With an urgent need to address this public health crisis, significant efforts have been made to develop pharmacological interventions such as liraglutide, semaglutide, and tirzepatide.^3–7^ These FDA-approved drugs exert their regulatory effect on appetite, body weight, and glucose homeostasis by targeting glucagon-like peptide 1 receptors (GLP-1Rs) expressed in peripheral tissues as well as the brain.^5, 8, 9^ GLP-1R agonists increase insulin release and decrease appetite leading to weight loss; however, with adverse side effects like nausea and vomiting, development of improved treatments for obesity are needed.^10–12^ Despite the widespread use of GLP-1R agonists as diabetes and obesity treatments, the role of the *endogenous* GLP-1 system, including neurons that produce GLP-1 in the brain, in mediating feeding and motivated behavior is unclear.

Feeding behavior is controlled by the central nervous system through the activation and inhibition of topographically organized and interconnected neural circuits sharing close three-dimensional space. Heroic efforts in basic research using rodent models has identified hypothalamic circuits that are critical for encoding hunger^13–18^, limbic circuits that are important for reward-based feeding^19–24^, and ascending hindbrain and brain stem circuits required for satiation and meal termination.^25–32^ Located in the rostral and caudal medulla, the nucleus of the solitary tract (NTS) is the first relay site from the glutamatergic vagus nerve receiving direct interoceptive signals from peripheral tissues including the stomach and intestines.^33–36^ The NTS is a complex structure comprised of heterogeneous groups of neurons that play a critical role in regulating satiety. Among these NTS neuronal populations are those that express preproglucagon (*Gcg* in mice) a precursor peptide that is processed to produce glucagon and GLP-1. These neurons, *Gcg*^NTS^ neurons (aka PPG neurons), have been identified as **the** brain’s primary source of glucagon-like peptide 1 (GLP-1).^37–40^

Numerous Fos mapping or imaging studies have shown that *Gcg*^NTS^ neurons are activated by gastric distension, gut satiety signals, large volume meals, and stressful/aversive stimuli.^38, 41–45^ These data, as well as anatomical studies demonstrating connections with the canonical hypothalamic-pituitary stress axis, suggest that *Gcg*^NTS^ neurons are important neuroendocrine regulators of behavior including feeding and stress coping.^46^ More recently, the development of transgenic animals providing targeted, high-fidelity genetic access to *Gcg*^NTS^ neurons has kindled a new wave of anatomical, physiological, and behavioral studies. These data illustrate that *Gcg*^NTS^ neurons make ascending projections throughout the brain and innervate hypothalamic, midbrain, and limbic nuclei.^37, 47, 48^ Studies examining the neurophysiology of *Gcg*^NTS^ neurons demonstrate that these neurons are spontaneously active and that the magnitude of their activity is enhanced by the application of circulating neuropeptidergic hormones like CCK, leptin, and oxytocin; and neuromodulators like serotonin and norepinephrine.^49–53^ Behavioral and physiological studies using chemogenetics have shown that *Gcg*^NTS^ neuron activation decreases feeding, while inhibition or ablation increases feeding during the consumption of large meals.^31, 38, 39, 52^ Together these studies provide a wealth of data pointing to the importance of *Gcg*^NTS^ neurons as *the endogenous source* of GLP-1 in the brain that sends axonal inputs to GLP-1R-expressing neurons in many brain sites.^54, 55^ However, the causal role of and modulation of *Gcg*^NTS^ neurons by feeding and energy status is unclear. Moreover, the lack of temporal and tunable activity using chemogenetics limits the ability to connect the specific activity states of *Gcg*^NTS^ neurons to their impact on feeding, motivation, and affect. In this study, we used anatomical, electrophysiological, and optogenetic approaches to answer these questions in both male and female mice. We observed that changes in energy status exert a major influence on the firing and excitability of *Gcg*^NTS^ neurons and that activation of *Gcg*^NTS^ neurons can produce sex-dependent changes in anxiety, avoidance, eating, and appetitive behavior.

## METHODS

### Animals

We used 8-16 week old C57BL6/J, *Gcg*-iCre (MGI: 5898434^56^, or *Dbh*-Cre (MGI: 6305870^57^) male and female mice throughout this study (Jackson Labs, Bar Harbor, ME). Additionally, we crossed *Gcg*-iCre mice with *Rosa26*-flx-stop-flx-L10-EGFP (MGI: 5548082^17^) reporter mice to generate *Gcg*-iCre:L10-eGFP (*Gcg*-GFP) male and female mice which were also used in this study. Unless otherwise specified, mice were group housed with 3-5 mice per cage on standard bedding and provided with food and water *ad libitum*. A 12:12 light cycle was maintained with lights on at 7AM and off at 7PM. All procedures were performed with approval from the University of Alabama at Birmingham Institutional Care and Use Committee (IACUC).

### Electrophysiology

#### Slice preparation

Electrophysiological experiments were performed as previously described.^20, 58^ Briefly, animals were removed from their cage and brought to the lab for brain slice preparation. The animal rested in a quiet chamber for 30 min prior to slice preparation to dissipate stress associated with animal transport from the vivarium. Mice were treated with a lethal dose of tribromoethanol (250 mg/kg, i.p.), and, after a deep plane of anesthesia was reached, animals were transcardially perfused with cold, sodium free N-methyl-D-glucamine (NMDG) artificial cerebrospinal fluid (aCSF)[(in mM) 93 N-methyl-D-glucamine, 2.5 KCl, 1.2 NaH_2_PO_4_, 30 NaHCO_3_, 20 HEPES, 25 Glucose, 5 L-ascorbic acid, 2 Thiourea, 3 sodium pyruvate, 10 MgSO_4_ X 7H_2_O, 0.5 CaCl_2_ X 2H_2_O]. All solutions were saturated with 95% CO_2_ and 5% O_2_. The brain was rapidly dissected and coronal 250 µM sections prepared in ice cold, oxygenated NMDG aCSF using a Leica VT1200S at 0.07 mm/s. Slices were immediately transferred to 36°C NMDG aCSF for 10 min, and then normal 36°C aCSF [(in mM): 124 NaCl, 4.4 KCl, 2 CaCl_2_, 1.2 MgSO_4_, 1 NaH_2_PO_4_, 10.0 glucose, and 26.0 NaHCO_3_]. Slices rested in normal aCSF for at least 30 min prior to recordings.

#### Recordings

Whole cell or cell-attached patch clamp recordings were performed in the NTS guided by DIC microscopy and GFP/tdTomato fluorescence. Slices were then transferred to a recording chamber (Warner Instruments), submerged in normal, oxygenated aCSF and maintained at 32°C with a flow rate of 2 ml/min. We patched GFP-expressing neurons in a balanced fashion in NTS slices. *Gcg*-GFP neurons were concentrated in the caudal NTS-containing sections surrounding the central canal or in the ventromedial portion of the dorsal vagal complex in sections containing a not fully formed area prostrema. For all recordings we used a potassium gluconate internal solution [(in mM): 135 C_6_H_11_KO_7_, 5 NaCl, 2 MgCl_2_, 20 HEPES, 0.6 EGTA, 4 Na_2_ATP, 0.4 Na_2_GTP at a final osmolarity of 290 mOsm at a pH of 7.3]. For voltage clamp recordings, neurons were voltage clamped at −70 mV using a Multiclamp 700B and currents were digitized with an Axon 1550B digitizer (Molecular Devices, Fremont, CA). For current clamp experiments, we recorded the excitability of *Gcg*^NTS^ neurons at their resting membrane potential and after injecting enough hyperpolarizing current so that their average potential was −70 mV (−15 to −45 pA). For optogenetic studies of ChrimsonR of *Gcg*^NTS^ neurons, 624 nM red-orange light was delivered through a 40X water immersion objective to neurons current clamped at −70 mV. A train of 60 pulses (1, 2.5, or 5 ms) were delivered to neurons at 1, 5, 10, 15, 20, and 40 Hz for three sweeps with an intersweep interval of ten seconds.

#### Data analysis

Data were analyzed in Clampfit 11.1 (Molecular Devices, San Jose, CA). Membrane capacitance and resistance were determined online using a −10 mV square pulse after the cell stabilized (∼1 min). We did not correct for liquid junction potential. Resting membrane potential was determined using a 2’ gap free current clamp recording where we took the average potential following a stabilizing period (usually 30’’). For rheobase, we identified the injected current at the peak of the first action potential. For recordings at −70 mV, we injected a steady variable amount of negative current using the amplifier while recording gap-free in current clamp until the cell reached −70 mV. For cell-attached analysis we used a threshold-based event detection to identify action-potential mediated currents across a 2-3 minute gap free recording for each neuron. Neurons had to demonstrate at least one identifiable action potential with appropriate kinetics to confirm the quality of the recording and loose seal.

### Stereotaxic Surgery

Surgical procedures were performed on 6-8 week old male and female *Gcg*-iCre mice. Animals were administered Buprenorphine SR (1.0 mg/kg, sc) and Meloxicam SR (5 mg/kg, sc) for analgesia directly prior to surgery and again on the second day of recovery. Mice were anesthetized under isoflurane inhalation (0.5 −5%) and placed in a stereotaxic frame. Eye lubricant was applied directly after the mouse was placed in the stereotaxic frame and reapplied as needed throughout surgery. Respiratory rate was visually monitored throughout surgical procedures. Internal temperature was monitored and maintained at 35°C using a homoeothermic biodynamic feedback system (Harvard Apparatus; Holliston, MA). After appropriate anesthetic depth was verified by a loss of toe pinch reflex, the scalp was manually depilated and sterilized via rotating application of 70% EtOH and betadine. We applied triple antibiotic ointment in addition to a topical lidocaine (4%) ointment for local analgesia. A midline incision was made, and the scalp parted to expose the skull surface. After verifying appropriate orientation of the head, 1 mm burr holes were drilled at the site of injection and fiber placement and in three additional locations for stability screw placement. For viral injections, a glass capillary was inserted into a Nanoject III (Drummond Scientific; Broomall, PA) and attached to the stereotaxic frame. Either AAV-FLEX-tdTomato (200nl, Addgene – cat# 28306-AAV5, lot# v122060) or pAAV-Syn-Flex-ro[ChrimsonR-tdTomato] (200nl, Addgene – cat# 62723-AAV5, lot# v63071), were injected at +/- 0.4mm lateral, −0.3mm posterior, z: −4.0mm ventral of obex. The rate per injection was 10 nl for 20 cycles, at a rate of 1 nl/s with a 30 second delay between each cycle. Optical fibers (black ceramic ferrule, 200 µM diameter, NA = 0.37, Hangzhou Braintech Technology; Hangzhou City, China) were secured in a custom-made ferrule holder (https://github.com/OpenBehavior/Modular-Stereotaxic-Holders) and implanted at a 15° angle +/- 1.2 mm lateral, - 0.3 mm posterior, −3.9 mm ventral of the obex. Set screws were shallowly fastened to the skull and a layer of Metabond was applied over the skull surface covering the base of the fibers and the screws. When the Metabond was dry, dental cement mixed with charcoal powder (to prevent laser light leakage and nonspecific behavioral effects) was applied.^59^ The mice were singly housed and remained in the colony for 3 weeks for recovery and to allow accumulation and expression of viral transgenes. Post-operative care was provided and body weight was monitored daily for 5 days or until mice achieved body weight within 5% of their initial preoperative weights.

### Animal Behavior

All behaviors were performed during the lights-on period in a cabinet at a light level of 25 lux. When not in the cabinet, animals remained in their home cages.

#### Open Field

Mice were habituated to handling and tethering to optical fiber patch cables (Doric Lenses; Quebec, Canada). The mice were tethered to optical patch cables and then allowed to explore the arena while providing 635 nm laser stimulation (Laser Century Inc. Shanghai, China) with a continuous 5 sec ON/OFF at 5 ms pulses to avoid heating effects. The frequency of the stimulation was 20 Hz, while the power of the laser was at 6 mW. Frequency and pulsewidth were controlled via an Arduino microcontroller. The overall time of each trial was set to 30 minutes. The dimensions for the arena were 50.165 × 50.165 cm. The arena was properly cleaned after each trial with water and the end of the day with dilute Blue Dawn soapy water. Data were analyzed using Noldus Ethovision XT version 16.

#### Real-time Place Preference

Mice were tethered to optical patch cables and then placed in a two-chamber arena, which was covered with fresh bedding per trial. Laser stimulation was provided as previously described. The frequencies tested were 1, 5, 10, 20 and 40 Hz. Trials lasted for a total of 30 minutes. Prior to each trial the stimulation side was reversed from the previous session so that real-time learning was always assessed. A separate reversal session occurred at 40 Hz for a total of 40 minutes where the side of stimulation was reversed at the 20-minute mark. The stimulation side was counterbalanced across both groups of animals throughout the entirety of the experiments. The boxes were thoroughly cleaned at the end of each day. Data were analyzed using Noldus Ethovision XT version 16.

#### Refeed

In preparation for this experiment animals were trained in a clear-walled box that allowed for access to a <u>F</u>eeding <u>E</u>xperimental <u>D</u>evice (FED).^60, 61^ Mice were trained to refeed from the FED for three one-hour sessions, while patch chords remained attached to the headcap. For experimental sessions, mice were then food deprived 24 hours prior to experiment. They were then placed in the arena with the FED. Each trial was 1 hr in length and stimulated using a 5 min OFF/ON epoch. Each ON period had a 5 sec ON/OFF interval for the laser to prevent heating. Data were analyzed using custom Python workflows to assign each pellet event to laser epochs.

#### FR1

Animals were food restricted (using ∼2g chow/day) during the period of FR1 training and testing. The same stimulation paradigm for refeed was used for the fixed ratio trials with the exception that the stimulation frequencies used were only 5 and 20 Hz. FEDs recorded the nose pokes of both the active and inactive side of the FED, while also dispensing a chocolate sucrose pellet only when the animal triggered the active port.

### *In vivo* Optogenetic Stimulation of *Gcg*^NTS^ Neurons – *Fos* validation

On trial day, individually housed mice were brought into the behavioral room and acclimated for 60 min prior to laser stimulation. Mice were tethered to bilateral fiber optic patch cords coupled to a 635 nm laser and placed in the home cage. After a 15-minute habituation period, laser stimulation was delivered as previously described at either 5 or 20 Hz for 5 min. Mice were then detached from patch cables and placed back in their home cage. Ninety minutes after the end of laser stimulation, mice were sacrificed with a terminal dose of tribromoethanol (250 mg/kg) and transcardially perfused with 50 ml of 0.01 M PBS followed by 50 ml 4% paraformaldehyde in 0.01 M PBS. Brains were extracted and post-fixed in 4% paraformaldehyde/PBS overnight then transferred to 30% sucrose for cryoprotection for 48-hours prior to cryosectioning and preserved for *Fos* immunostaining.

### Intermittent Access HFD consumption paradigm

*Gcg*-GFP male (n = 11) and female (n = 13) mice were individually housed 1 week prior to experimentation. We adapted our previously published method for these studies.^62^ Throughout the 5-day intermittent access period, mice were maintained with *ad libitum* access to standard chow and water. During the first 4 days, all mice were allowed *ad libitum* access to a high fat diet (HFD) containing 60% fat (D12492; Research Diets, Inc., New Brunswick, NJ) for 1 hr per day at ∼11 AM. Each day, bodyweight measurements were taken and the amount of HFD food intake was measured for each mouse. On day 5, mice were allocated to one of the following groups: *ad libitum* (AL), 50% *ad libitum* (AL50), 30% *ad libitum* (AL30), or food deprived (FD). Mice in the AL group (n = 6) were provided *ad libitum* access to HFD for 1 hr. For groups AL50 (n = 6) and AL30 (n = 6), the average amount of HFD consumed on day 4 was calculated and 50% or 30% of that amount respectively was provided for 1 hr on day 5. Mice in the AL30 and AL50 groups always consumed the provided HFD. Mice allocated to the FD group (n = 6) were not given HFD access on day 5. Following the 1 hr intermittent access period on day 5, bodyweights and food consumption were measured, and a 90 min timer was started to allow for *Fos* protein expression. After 90 min, mice were perfused as previously described and the brain extracted.

### Histology

A lethal dose of tribromoethanol (250 mg/kg) was administered via intraperitoneal injection prior to sacrifice. Animals were then transcardially perfused after appropriate depth of anesthesia was confirmed by a loss of the toe pinch reflex. Transcardial perfusions were performed as before. Brains were extracted, post-fixed in 4% paraformaldehyde overnight then transferred to 30% sucrose for 36-48 hr. Once sufficiently cryoprotected, brains were cryosectioned at 40 µm. Horizontal tissue sections were directly mounted to Superfrost plus microscope slides (Fisher Scientific; Waltham, MA). Coronal sections were either directly mounted to Superfrost plus microscope slides (Fisher Scientific) or collected in 1:1 glycerol/PBS and mounted to Superfrost microscope slides (Fisher Scientific). Slides were coverslipped with hardset mounting media containing DAPI (Vector Laboratories; Newark, CA).

#### Fluorescence in situ hybridization (FISH)

Fresh brain tissue was harvested from 8-16 week old C57BL6/J or *Gcg*-iCre male and female mice and flash frozen on dry ice then placed in an air-tight container and stored at −80°C. At least 1 hour prior to cryosectioning, tissues were moved from −80°C into the cryostat (−20°C) to allow for temperature acclimation. Coronal sections (18 µm) were collected and directly mounted onto Superfrost Plus slides (Fisher Scientific). Slides were stored in a light-tight slide box, sealed with CryoHUG™ tape (Fisher Scientific) to prevent moisture accumulation, and stored at −80°C. Single molecule RNAscope® fluorescent multiplex assays were performed in accordance with the manufacturer’s protocol (Advanced Cell Diagnostics; Newark, CA). For validation of the *Gcg*-iCre mouse line, targets probes for *Gcg* (Mm-Gcg-C2, Cat No. 400601-C2, Accession No. NM_008100.3) and *iCre* (iCre-C3, Cat No. 423321-C3, Accession No. AY056050.1) were used for the RNAscope® multiplex fluorescent assay. For assessment of *Dbh* and *Gcg* mRNA expression patterns in C57BL6/J mice, target probes for *Dbh* (Mm-Dbh, Cat No. 407851, Accession No. NM_138942.3) and *Gcg* (Mm-Gcg-C2, Cat No. 400601-C2, Accession No. NM_008100.3) were used. Appropriate negative controls were included in each assay to verify probe specificity.

#### Fluorescence Immunohistochemistry

*Fos* immunohistochemistry was performed as previously described with some modifications.^20^ Free-floating (40 µm) coronal tissue sections were washed twice in 0.01 M PBS for 10 min, then permeabilized for 30 min in 50% methanol (diluted in ddH_2_O). Endogenous peroxidases were quenched with a 5 min incubation in 3% hydrogen peroxide. Sections were then washed twice in 0.01 M PBS for 10 min followed by a 1 hr incubation in a blocking solution comprised of 0.3% Triton X-100 (Fisher Scientific) and 1% Bovine Serum Albumin (BSA; Sigma, St. Louis, MO). Tissue sections were then incubated in the blocking solution containing the guinea pig anti-c-Fos primary antibody (1:1000; Cat No. 226-308, Synaptic Systems; Gottingen, Germany) overnight at 4°C with constant agitation. Sections were then washed in TNT buffer comprised of dH2O containing 0.15 M NaCl (Fisher Scientific), 0.1 M Tris HCl (pH 7.5, Fisher Scientific), and 0.05% Tween 20 (Fisher Scientific) for 10 min. Sections were then incubated in TNB blocking buffer comprised of dH2O containing 0.1 M Tris-HCl (pH 7.5, Fisher Scientific), 0.15 M NaCl (Fisher Scientific), and 0.5% blocking reagent (Cat No. K1052; APExBIO, Houston, TX) for 30 min. Sections were then incubated for 30 min in a Horse Radish Peroxidase conjugated goat anti-guinea pig secondary antibody (1:100; Cat No. 106-035-003, Jackson ImmunoResearch; West Grove, Pennsylvania) diluted in TNB buffer, washed four times in TNT buffer for 5 min/wash, then incubated in Cyanine 5 Tyramide (1:200, APExBIO) diluted in TSA amplification diluents (APExBIO) for 10 min followed by two washes in TNT buffer for 10 min. Sections were then mounted on SuperFrost slides (Fisher Scientific), allowed to dry, then coverslipped using hardset mounting medium with DAPI (Vector Labs).

### Imaging and Image Analysis

Horizontal tissue sections obtained from *Gcg*-GFP male (n = 3) and female mice (n = 5) were imaged using a Keyence BZ-X800 fluorescence microscope (Keyence; Itasca, IL) under 10x magnification with 2-4 tissue sections imaged per mouse. Cell counts were performed using object-based segmentation modules in CellProfiler™ 4 (Broad Institute of MIT and Harvard; Cambridge, MA). Coronal sections obtained from *Gcg*-GFP male (n = 11) and female (n = 13) mice were imaged using the Keyence BZ-X800 fluorescence microscope under 20x magnification. *Gcg* expressing neurons were counted using the cell counter tool in ImageJ/Fiji (NIH). An unpaired, two-tailed Student’s *t* test was performed in GraphPad Prism 9 for data obtained from horizontal sections as well as data obtained from coronal sections.

For all animals used in optogenetics experiments, 40 µm coronal sections were obtained across the NTS and inspected using an Olympus BX43 microscope with epifluorescence optics to verify fiber placement and virus expression. Representative images were captured using a Keyence BZ-X800 fluorescence microscope under 20x magnification. Optical fiber placement was determined by comparison to a mouse brain atlas (Paxinos and Watson). Mice not showing viral expression or failed fiber placement were excluded from subsequent data analysis.

#### Quantification of FISH labeling

For all coronal tissue sections used for *in situ* hybridization assays, tiled z-stacks of the NTS and surrounding regions were captured using a Keyence BZ-800 fluorescence microscope under 40x magnification. Stitched, maximum intensity projections of raw images were obtained using the Keyence Analyzer software. Resulting split channel images were converted to 16-bit in ImageJ/Fiji (NIH). Prior to background subtraction, nonspecific background signal was removed using image subtraction where 488 nm generated images were subtracted from 550 nm and 647 nm images. Composite images were generated, and the region of interest (NTS) defined using the freehand selection tool in ImageJ. Area measurements were obtained for each defined region and cells counted using the cell counter tool in ImageJ. For analysis of *Gcg* and *iCre* mRNA coexpression, *Gcg* mRNA was labeled with Cy3 (550 nm excitation) and *iCre* mRNA was labeled with Cy5 (647 nm excitation). Cell counts were obtained for *Gcg* mRNA-positive cells, *iCre* mRNA-positive cells, and *Gcg*/*iCre* mRNA-positive cells. For analysis of *Dbh* and *Gcg* mRNA expression, *Dbh* mRNA was labeled with Cy3 (550 nm excitation) and *Gcg* mRNA was labeled with Cy5 (647 nm excitation). Cell counts were obtained for *Dbh* mRNA-positive cells, *Gcg* mRNA-positive cells, and *Dbh*/*Gcg* mRNA-positive cells.

#### Quantification of Fos expression

Fluorescence immunohistochemistry images were taken using a Keyence BZ-X800 fluorescence microscope under 20x magnification. Tiled z-stacks were captured of the NTS and surrounding region. Stitched, maximum intensity projections of raw images were obtained using the Keyence Analyzer software. Resulting split channel images were converted to 16-bit in ImageJ. Background subtraction was employed to reduce noise and composite images generated. Using the freehand selection tool, the NTS was defined and area measurements taken. *Gcg+* neurons were counted in addition to neurons that co-expressed *Gcg* and *Fos* using the cell counter tool in ImageJ. To obtain overall *Fos*+ cell counts, object-based segmentation was used in Cell Profiler where nuclei (DAPI) and *Fos* signals were segmented as individual primary objects. Cells expressing *Fos* were identified using object filtering where only counts for nuclei objects overlapping with *Fos* objects were included in counts. Accuracy of object segmentation was verified for each image to ensure false positives were not included in overall counts. Inclusion criteria for *Fos* analysis required each section to contain *Gcg*+ cell bodies. The average number of *Gcg*+, *Fos*+, and *Gcg*+/*Fos*+ cells was calculated per animal for further analysis.

For the intermittent access HFD experiment, *Fos* analysis was performed using coronal sections obtained from *Gcg*-GFP male (n = 11) and female (n = 13) mice where 2-4 sections were imaged and analyzed per mouse. The average proportion of *Gcg*+ cells expressing *Fos* was calculated as an assessment of *Gcg* cells. A one-way ANOVA was used to determine significant differences among feeding group followed by Dunnett’s multiple comparisons analysis with each group compared to the control group (no HFD exposure on day 5). For the *ad libitum* group (AL), a Pearson’s correlation analysis was performed to assess the relationship between meal size (expressed as % bodyweight) and the average proportion of *Fos* expressing *Gcg*^NTS^ cells.

The optogenetic experiment used male *Gcg*-iCre mice (n = 12) with 2-4 images analyzed per mouse. The average proportion of *Gcg*+ cells expressing *Fos* was calculated as an assessment of *Gcg*^NTS^ cells. Additionally, the average density (expressed as cells per mm^2^) of non-*Gcg*^NTS^ cells expressing *Fos* was assessed. For both measurements, a one-way ANOVA was performed followed by Tukey’s multiple comparisons test.

### Statistics

Statistical analyses were performed using GraphPad Prism 9.5.1. Data presented in graphs are expressed as the mean ± standard error of the mean (s.e.). Significant differences were determined using a *p*-value < 0.05. Figures were assembled in Adobe Illustrator.

## RESULTS

### Anatomy of Gcg^NTS^ Neurons and Mouse Line Validation

We generated male and female *Gcg-*iCre; *Rosa26-*flx-stop-flx-L10-EGFP (*Gcg*-GFP mice) mice and used microscopy to count the number and distribution of *Gcg*^NTS^ neurons in adult *Gcg*-GFP mice. GFP-labeled *Gcg* neurons were located in the caudal and ventrolateral NTS, which is consistent with previous findings in both rats and mice.^37, 63–65^ We counted *Gcg*^NTS^ neurons in coronal (**FIG. 1A, B**) and horizontal (**FIG. 1C**) fixed tissue sections from male and female mice. Results from these analyses revealed no significant differences in *Gcg*^NTS^ density (cells/mm^2^) between male and female *Gcg*-GFP mice (**FIG. 1B, C**). To better understand our published *Gcg-* iCre mouse line^56^; we performed fluorescence *in situ* hybridization and quantified *Gcg* and *iCre* mRNA in the NTS (**FIG. 1D-I)**. We found that *iCre* was expressed in ∼70% of *Gcg* neurons (**FIG. 1H**) and 94% of *iCre*-expressing cells also expressed *Gcg* (**FIG. 1I**). Therefore, we established that our *Gcg-iCre* transgenic provides acceptable penetrance and high-fidelity genetic access to *Gcg*^NTS^ neurons.

**Figure 1.**
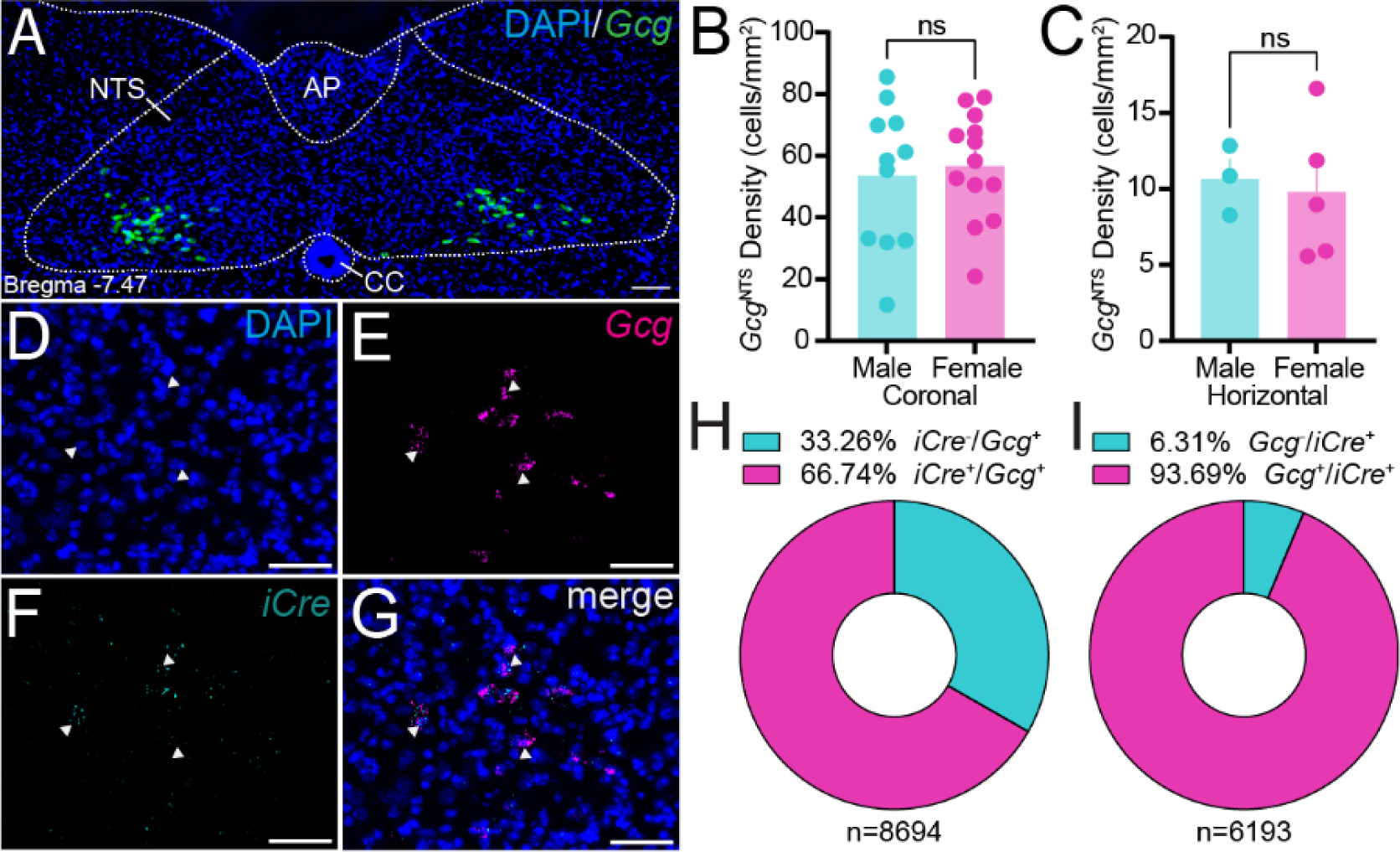
Anatomical distribution of *Gcg*^NTS^ cells and validation of *Gcg*-iCre mouse line. (**A**) Representative image of coronal tissue section from a *Gcg*-GFP mouse showing *Gcg*^NTS^ neurons (green). Scale bar = 500µm. (**B**) Quantification of *Gcg*^NTS^ neurons per mm^2^ obtained from coronal tissue sections from *Gcg*-GFP mice (n = 24; 11 male, 13 female) with 2-4 sections/mouse. Unpaired Student’s *t* test; F (10, 12) = 1.799, *p* = 0.708 (male vs. female). (**C**) Quantification of *Gcg*^NTS^ neurons per mm^2^ obtained from horizontal tissue sections from *Gcg*-GFP mice (n = 8; 3 male, 5 female) with 2 sections/mouse. Unpaired Student’s *t*-test; F (4, 2) = 4.011, *p* = 0.7748 (male vs. female). (**D-G**) Single channel enlarged view of NTS neuron showing (**D**) DAPI (blue), (**E**) *Gcg* mRNA expression (magenta), (**F**) *iCre* mRNA expression (cyan), and (**G**) a merged view. White arrows indicate neurons coexpressing *Gcg* and *iCre* mRNA. Scale bar = 50µm. (**H**) Quantification of *iCre* mRNA expression within *Gcg*+ cells (n = 8694 cells). (**I**) Quantification of *Gcg* mRNA expression within *iCre*+ cells (n = 6193 cells). For **H-I**, data were obtained from 13 images taken from *Gcg-*iCre mice (n = 3) with 2-6 images/mouse. For **H**, the mean number of *Gcg*+ cells counted per image was 669 ± 98.8 (s.e.). For **I**, the mean number of *iCre*+ cells counted per image was 476 ± 78.5 (s.e.).

Anatomical data from male rats and mice demonstrate that *Gcg*^NTS^ neurons are distinct from larger clusters of neurons comprising the A2 dorsal vagal complex that produce the norepinephrine and express the enzyme dopamine β-hydroxylase (*Dbh*).^49, 63^ To confirm this observation in male and female C57BL6/J mice, we used fluorescence *in situ* hybridization and quantified the presence and coexpression of *Gcg* and *Dbh* mRNA in the NTS (**Supplemental FIG. 1**). We observed little overlap in these mRNAs where *Gcg* mRNA was expressed in ∼8% of *Dbh* mRNA+ cells (**SUP. FIG. 1E**) and *Dbh* mRNA was expressed in 14% of *Gcg* mRNA+ cells (**SUP. FIG. 1F**). We did not observe any difference in the density and number of *Gcg* and *Dbh* mRNA+ cells between males and female mice (data not shown).

### Impact of Energy and Feeding State on Gcg^NTS^ Neuron Activation In Vivo

To assess the activation of *Gcg*^NTS^ neurons in response to food intake we adapted a previously used paradigm in rats.^41, 42^ We provided male and female *Gcg*-GFP mice with intermittent access to a high fat diet (HFD) in their home cage for 1 hour per day for four days to generate entrained patterns of large volume food intake (**FIG. 2A**). On the fifth day, mice were randomly assigned to one of the following groups: *ad libitum* (AL), 50% *ad libitum* (AL50), 30% *ad libitum* (AL30), or food deprived (FD). Intermittent access to HFD was then provided according to group assignment for 1 hour after which the HFD was removed (**FIG. 2A**). As expected, the average amount of HFD consumed increased over the four-day course (**FIG. 2B**); however, a slight decrease in bodyweight was observed across all animals over the five-day course (**FIG. 2C**). Analysis of *Fos* expression in *Gcg*^NTS^ neurons revealed no significant differences in the proportion of *Fos*+ *Gcg*^NTS^ among feeding groups with respect to the FD control group. AL mice demonstrated a marginal increase in the proportion of *Fos* expression within *Gcg*^NTS^ cells that did not reach statistical significance (**FIG. 2D**). Due to the intrinsic variability of HFD consumption for AL mice on day five, we examined the correlation between the proportion of *Fos* expressing *Gcg*^NTS^ cells and meal size (expressed as percent body weight). Results from this analysis indicated a significant positive correlation between food intake and *Fos+ Gcg*^NTS^ cells (**FIG. 2E**). While AL mice demonstrated the greatest proportion of *Fos* expression among *Gcg*^NTS^ cells (**FIG. 2D**), histological examination revealed *Fos* expression in an alternative NTS neuronal subpopulation that may be modulated by feeding state (**FIG. 2F-I**).

**Figure 2.**
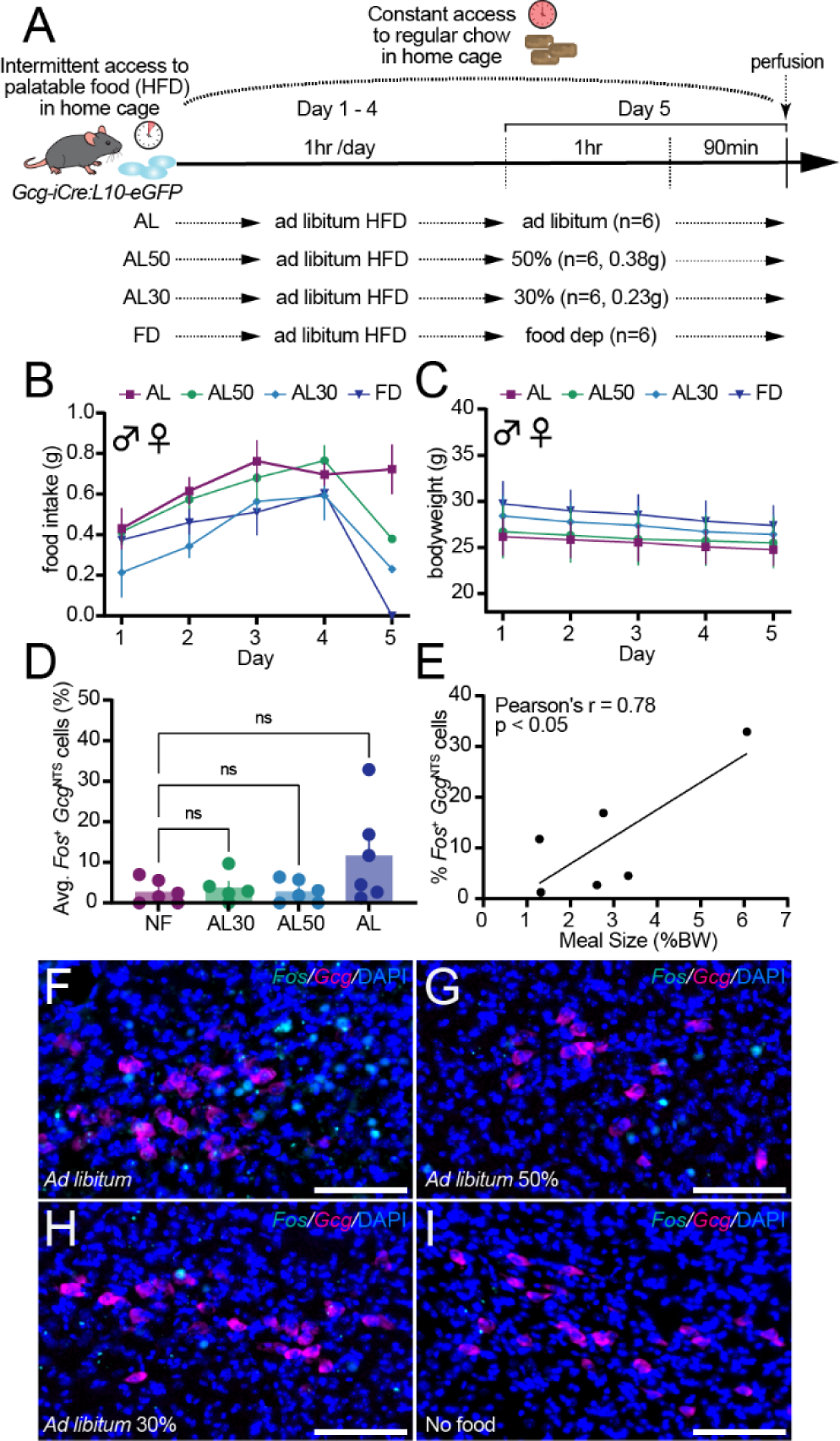
*Gcg*^NTS^ neuron activation in response to intermittent high fat diet consumption. (**A**) Schematic of experimental paradigm to provide intermittent access to energy-dense, palatable food for 5 days. (**B**) Food intake across 5 days with combined sexes. (**C**) Body weights across 5 days with combined sexes. For B-C, 24 *Gcg*-GFP male (n = 12) and female (n = 12) mice were used. (**D**) Quantification of the proportion of *Gcg*^NTS^ cells that expressed *Fos* across groups with combined sexes. Each group consisted of 5-7 mice with 2-3 males/group and 3-4 females/group. For each animal, 2-4 images were analyzed. Data were analyzed using an ordinary one-way ANOVA with a Dunnett’s multiple comparisons test. F (3, 19) = 2.437, *p* = 0.0962. NF vs AL30: *p* = 0.9876, NF vs AL50: *p* > 0.9999, NF vs. AL: *p* = 0.0831. (**E**) Correlation of meal size (expressed as % bodyweight) and % *Gcg*^NTS^ cells that were *Fos*+ for *ad libitum* group only (n = 6; 3 male, 3 female). Pearson’s r = 0.7830, *p* = 0.0328. (**F-I**) Merged magnified representative images of *Gcg*^NTS^ cells and *Fos* for the (**F**) AL group, (**G**) AL50 group, (**H**) for AL30 group, and (**I**) NF group where *Gcg* expression is shown in magenta, *Fos* expression is shown in cyan, and DAPI is shown in blue. Pseudo color section intended for ease of visualization. For **F-I** scale bar = 100µm.

### Energy status alters physiology and excitability of Gcg^NTS^ neurons

Characterization of *Gcg*^NTS^ activity states via *Fos* mapping provides only a snapshot of the potential changes in neural physiology and is inherently bimodal. To gain a deeper understanding of how *Gcg*^NTS^ neuron physiology is altered by energetic status and feeding, we used *ex vivo* electrophysiology in *Gcg*-GFP mice (**FIG. 3A-D**). Mice were randomly assigned to three groups: *ad libitum* chow fed (AL), 24-hour food deprived (FD), or 24-hour food deprived and then chow refed (RF, **FIG. 3E**). RF animals ate 0.87±0.07 grams of chow with no significant differences between male and female *Gcg*-GFP mice, although a nonsignificant increase in female mice was noted (**FIG. 3F**). First, to understand how energy status impacts *Gcg*^NTS^ neuron firing without potentially dialyzing the intracellular contents of *Gcg*^NTS^ neurons, we used cell-attached recordings of *Gcg*^NTS^ neurons in our three-group design (**FIG. 3G-H**).^66^ Consistent with previous data using perforated-patch recordings^49^, we found that almost every *Gcg*^NTS^ neuron in the AL group displayed action-potential-mediated currents (**FIG. 3I**) and there was no significant difference in the mean firing rate of *Gcg*^NTS^ neurons between male and female mice (2.9±0.29 Hz vs 2.5±0.18 Hz). We observed a significant decrease in the mean firing rate of *Gcg*^NTS^ neurons in FD animals, and no significant difference in RF animals relative to AL controls (**FIG. 3I**). Interestingly, this phenomenon was specific to male *Gcg*-GFP mice (**FIG. 3J+K**). To assess this potential sex difference more directly, we compared the mean firing rate of male and female FD mice (1.3±0.23 Hz vs 2.8±0.32 Hz, *p* = 0.0003, not shown). Although we observed no difference in the mean firing rates between AL vs RF animals, we noted a much higher tail suggesting greater variance and higher firing rates of *Gcg*^NTS^ neurons in the RF condition. Remarkably, we observed individual *Gcg*^NTS^ neurons firing at 7-9 Hz in *ex vivo* slices. A separate analysis comparing the distribution of *Gcg*^NTS^ neuron firing rate between AL vs RF mice revealed a significant difference (**FIG. 3L**). These data demonstrate that food deprivation and refeeding persistently alters the firing rate of *Gcg*^NTS^ neurons.

**Figure 3.**
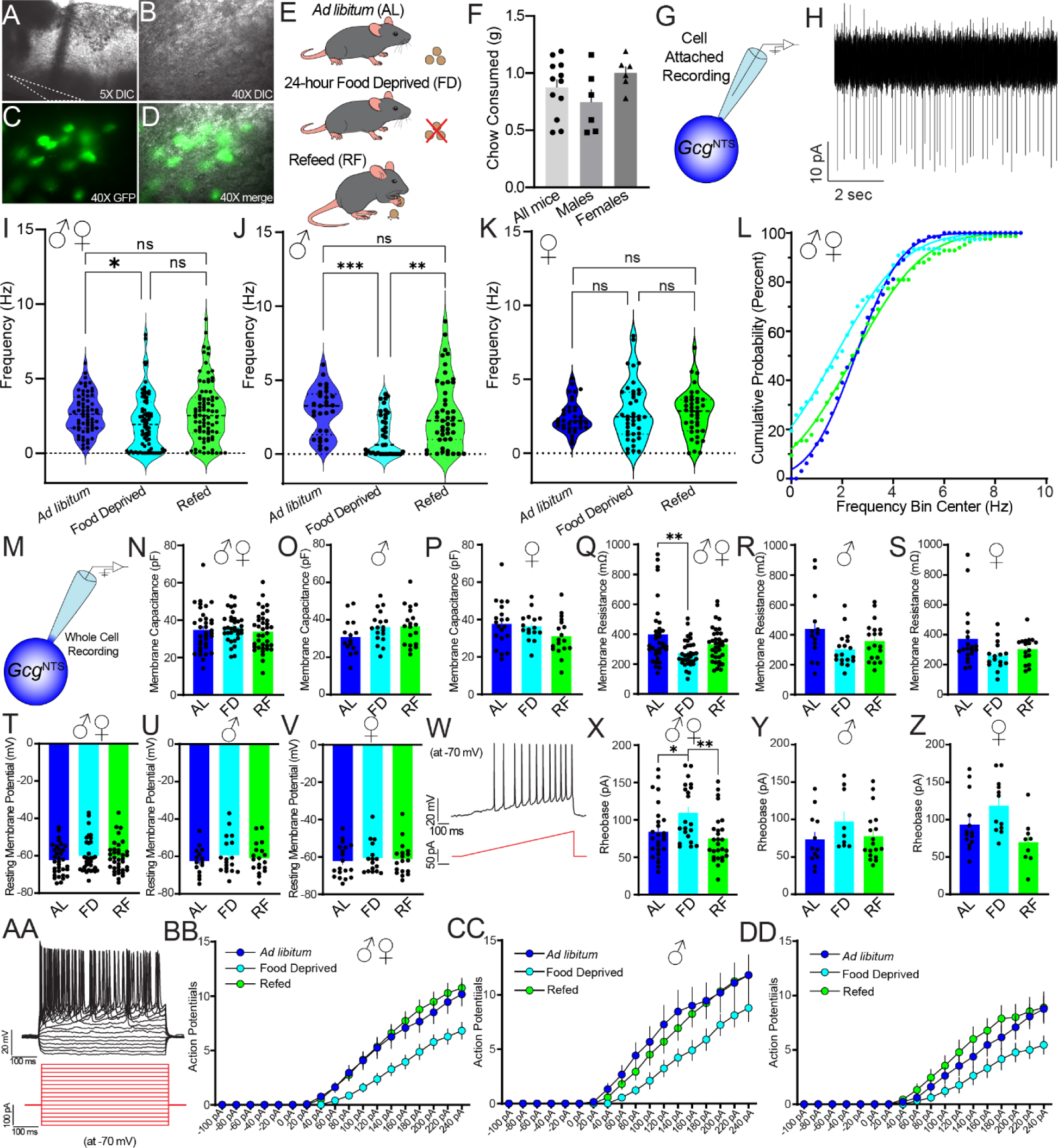
Modulation of *Gcg*^NTS^ neuron firing, intrinsic properties, and excitability by energy and feeding state. (**A**) 5X and (**B**) 40X DIC view of NTS containing brain stem slices. (**C-D**) GFP and merged view of GFP+ and putative *Gcg*^NTS^ neurons under 40X objective. (**E**) Schematic of three experimental groups for recording. *Ad libitum* (AL) = mice were given normal access to chow up to the day of slice preparation. Food Deprived (FD) = chow was removed ∼24 hours prior to slice preparation. Refeed (RF) = chow was removed ∼24 hours prior and then animals were given access to *ad libitum* chow for 1 hour immediately prior to slice preparation. (**F**) Chow consumed for Refeed mice recorded from in Figure 3. Data from male and female mice were analyzed using an unpaired Student’s *t* test, *p* = 0.0705. (**G**) Schematic of cell-attached or “loose-patch” recording used in data panels **H-L**. (**H**) Example trace of *Gcg*^NTS^ neuron action-potential mediated currents in cell-attached recordings. Average spike amplitudes = 66.781±4.94 pA (n = 229 cells across all groups). (**I**) Firing frequency of *Gcg*^NTS^ neurons across all AL, FD, and RF mice. AL vs FD: *p* = 0.0350, AL vs RF: *p* = 0.9999, FD vs RF: *p* = 0.0808 (n = 229 cells from 18 mice, 6/group) (**J**) Firing frequency of *Gcg*^NTS^ neurons across male AL, FD, and RF mice. AL vs FD: *p* = 0.0005, AL vs RF: *p* = 0.9128, FD vs RF: *p* = 0.0057 (n = 114 cells from 9 mice, 3/group). (**K**) Firing frequency of *Gcg*^NTS^ neurons across female AL, FD, and RF mice. AL vs FD: *p* > 0.9999, AL vs RF: *p* > 0.9999, FD vs RF: *p* > 0.9999 (n = 115 cells from 9 mice, 3/group). (**L**) Cumulative distribution of firing frequency of *Gcg*^NTS^ neurons across mice AL, FD, and RF mice. AL vs FD: *p* = 0.0282, AL vs RF: *p* = 0.0150, FD vs RF: *p* = 0.0001. (**M**) Schematic of whole-cell recording configuration used in data panels **N-DD**. (**N**) Membrane capacitance of *Gcg*^NTS^ neurons recorded from all mice. F (2, 103) = 0.4912, *p* = 0.6133. (**O**) Membrane capacitance of *Gcg*^NTS^ neurons recorded from male mice. F (2, 49) = 1.734, *p* = 0.1872. (**P**) Membrane capacitance of *Gcg*^NTS^ neurons recorded from female mice. F (2, 51) = 2.161, *p* = 0.1257. (**Q**) Membrane resistance of *Gcg*^NTS^ neurons recorded from all mice. F (2, 100) = 6.232, *p* = 0.0028. AL vs FD: *p* = 0.0019, AL vs RF: *p* = 0.1634, FD vs RF: *p* = 0.2658. (**R**) Membrane resistance of *Gcg*^NTS^ neurons recorded from male mice. F (2, 48) = 3.027, *p* = 0.0578. (**S**) Membrane resistance of *Gcg*^NTS^ neurons recorded from female mice. F (2, 50) = 2.869, *p* = 0.0662. (**T**) Resting membrane potential of *Gcg*^NTS^ neurons recorded from all mice. F (2, 102) = 0.6825, *p* = 0.5077. (**U**) Resting membrane potential of *Gcg*^NTS^ neurons recorded from male mice. F (2, 49) = 0.5123, *p* = 0.6023. (**V**) Resting membrane potential of *Gcg*^NTS^ neurons recorded from female mice. F (2, 50) = 0.1848, *p* = 0.8318. (**W**) Example trace of *Gcg*^NTS^ neuron current clamped with hyperpolarizing current to hold at −70 mV (black) and then injected with a depolarizing current ramp (red) to evoke action potentials. (**X**) Rheobase of *Gcg*^NTS^ neurons recorded from all mice. F (2, 71) = 5.657, *p* = 0.0053. AL vs FD: *p* = 0.0467, AL vs RF: *p* = 0.6447, FD vs RF: *p* = 0.0044. (**Y**). Rheobase of *Gcg*^NTS^ neurons recorded from male mice. F (2, 36) = 1.323, *p* = 0.2791. (**Z**) Rheobase of *Gcg*^NTS^ neurons recorded from female mice. F (2, 32) = 4.887, *p* = 0.0141. AL vs FD: *p* = 0.1851, AL vs RF: *p* = 0.2832, FD vs RF: *p* = 0.0107. (**AA**) Example trace of *Gcg*^NTS^ neuron in current clamped with hyperpolarizing current to hold at −70 mV (black) and then injected with increasing depolarizing current steps (red) to evoke action potentials. (**BB**) Current step evoked action potential number for *Gcg*^NTS^ neurons recorded from all mice. Feeding State X Current Step Interaction effect: F (34, 1166) = 3.923, *p* < 0.0001; Feeding effect: F (2, 69) = 5.118, *p* = 0.0085; Current step effect: F (1.601, 109.8) = 169.3, *p* < 0.0001. Significant decreases between AL/RF and FD between +80-240 pA. (**CC**) Current step evoked action potential number for *Gcg*^NTS^ neurons recorded from male mice. Feeding State X Current Step Interaction effect: F (34, 571) = 1.520, *p* < 0.0001; Feeding effect: F (2, 34) = 2.065, *p* = 0.1424; Current step effect: F (1.609, 54.05) = 90.92, *p* < 0.0001. No significant post-tests. (**DD**) Current step evoked action potential number for *Gcg*^NTS^ neurons recorded from female mice. Feeding State X Current Step Interaction effect: F (34, 544) = 2.380, *p* < 0.0001; F (2, 32) = 3.003, *p* = 0.0638; Current step effect: F (1.598, 51.12) = 83.60, *p* < 0.0001. No significant post-tests. For (**I-J**), data were analyzed using Kruskal-Willis test with Dunn’s multiple comparisons test. For **L**, cumulative distributions were analyzed using Kolmogorov-Smirnov test. For **N-Z**, data were analyzed using one-way ANOVA with Sidak’s multiple comparisons tests. For **BB-DD**, data were analyzed using two-way repeated measures ANOVA with Tukey’s multiple comparisons test.

To assess whether changes in energy status affect the intrinsic neurophysiology of *Gcg*^NTS^ neurons, we used whole cell patch clamp electrophysiology using *Gcg*-GFP mice (**FIG. 3M**). We observed no significant difference in the membrane capacitance (Cm) of *Gcg*^NTS^ neurons across AL, FD, and RF states overall or in either sex (**FIG. 3N-P**). Thus, there is no major difference in *Gcg*^NTS^ neuron’s ability to hold charge likely reflecting cell size across these energetic states. Interestingly, we observed a significant decrease in membrane resistance (Rm) in FD animals relative to AL animals (**FIG. 3Q**). Separate analyses in both male and female mice revealed a nonsignificant decrease between AL and FD animals (**FIG. 3R, S**). In all analyses, RF animals showed a higher Rm than FD animals, but this did not reach statistical significance. These data suggest that there is more conductance and open ion channels in *Gcg*^NTS^ neurons in a FD state. To determine if energy status affects *Gcg*^NTS^ neuron membrane potential, we recorded the resting membrane potential (RMP) in current clamp mode. We observed no significant difference in RMP as a function of energy status or sex (**FIG. 3T-V**). We next examined how energy status impacts the excitability of *Gcg*^NTS^ neurons in current-clamp mode. Since we observed a wide range of RMP of *Gcg*^NTS^ neurons, we performed these experiments at RMP (not shown) and when clamped with enough hyperpolarizing current to hold the neuron at −70 mV on average (**FIG. 3W**). We observed a significant increase in the amount of current needed to evoke an action potential (rheobase) in FD animals relative to AL animals and RF animals (**FIG. 3X**). Separate analyses by sex revealed similar nonsignificant trends in male mice (**FIG. 3Y**), and RF animals showed a significant decrease in rheobase relative to FD animals (**FIG. 3Z**). There was no significant difference in the rheobase of AL animals between male and female mice (data not shown). Last, we examined the ability of *Gcg*^NTS^ neurons to initiate and sustain volleys of action potentials in response to depolarizing current steps (500 ms, **FIG. 3AA**). We observed a dramatic reduction in current step-evoked action potentials in FD *Gcg*^NTS^ neurons that appeared as a rightward shift and maximal decrease in the number of action potentials generated at each current step (**FIG. 3BB**). In analyses separated by sex, we observed a significant difference in current step-evoked action potentials (**FIG. 3CC-DD**) in both male and female mice. When we directly compared current step-evoked action potentials in male and female AL animals, we observed a significant decrease in the number of evoked action potentials at each current step. Together these data demonstrate that food deprivation alters membrane resistance and decreases the excitability of *Gcg*^NTS^ neurons and that the excitability of *Gcg*^NTS^ neurons differs by sex.

### ChrimsonR allows for tunable and reversible activation of Gcg^NTS^ neurons in vivo and ex vivo

Using male *Gcg*-iCre mice we expressed the red light-gated opsin ChrimsonR or tdTomato in *Gcg*^NTS^ neurons and implanted angled bilateral optical fibers terminating dorsal of the brainstem, approximately ∼250 µM above *Gcg*^NTS^ neurons (**FIG. 4A**).^67^ After recovery from surgery, we assayed the activation of *Gcg*^NTS^ neurons after a brief 5 min photostimulation session delivered at 5 or 20 Hz in a home cage (**FIG. 4B**). To determine how tunable optogenetic stimulation is via ChrimsonR, ChrimsonR-expressing animals received stimulation at 5 or 20 Hz while TdTomato controls only received 20 Hz. Ninety minutes after stimulation, we perfused animals and performed immunostaining for *Fos* in NTS sections. Remarkably, 5-minute stimulation of *Gcg*^NTS^ neurons at 20 Hz in ChrimsonR-expressing animals induced the expression of *Fos* in ∼85% of ChrimsonR-expressing *Gcg*^NTS^ neurons, whereas 5 Hz only resulted in *Fos* expression in <40% of ChrimsonR-expressing *Gcg*^NTS^ neurons (**FIG. 4C**). Moreover, the *Fos* expression observed for both ChrimsonR stimulation groups (5 Hz and 20 Hz) was significantly greater than that observed for the control group (TdTomato 20 Hz) with <2% of *Gcg*^NTS^ cells expressing *Fos* (**FIG. 4C**). We also noted a nonsignificant increase in the density of *Fos* expression in non-*Gcg*^NTS^ cells for all groups (**FIG. 4D**). Histological assessment of NTS sections (**FIG. 4E-P**) revealed robust expression of TdTomato and ChrimsonR in putative *Gcg*^NTS^ neurons, and we could identify sites of optical fiber termination passing through the posterior cerebellum and dorsal parts of the medulla at the level of the gracile nucleus (not shown). These data establish that our light-gated opsin, ChrimsonR, is an effective strategy to activate *Gcg*^NTS^ neurons *in vivo*.

**Figure 4.**
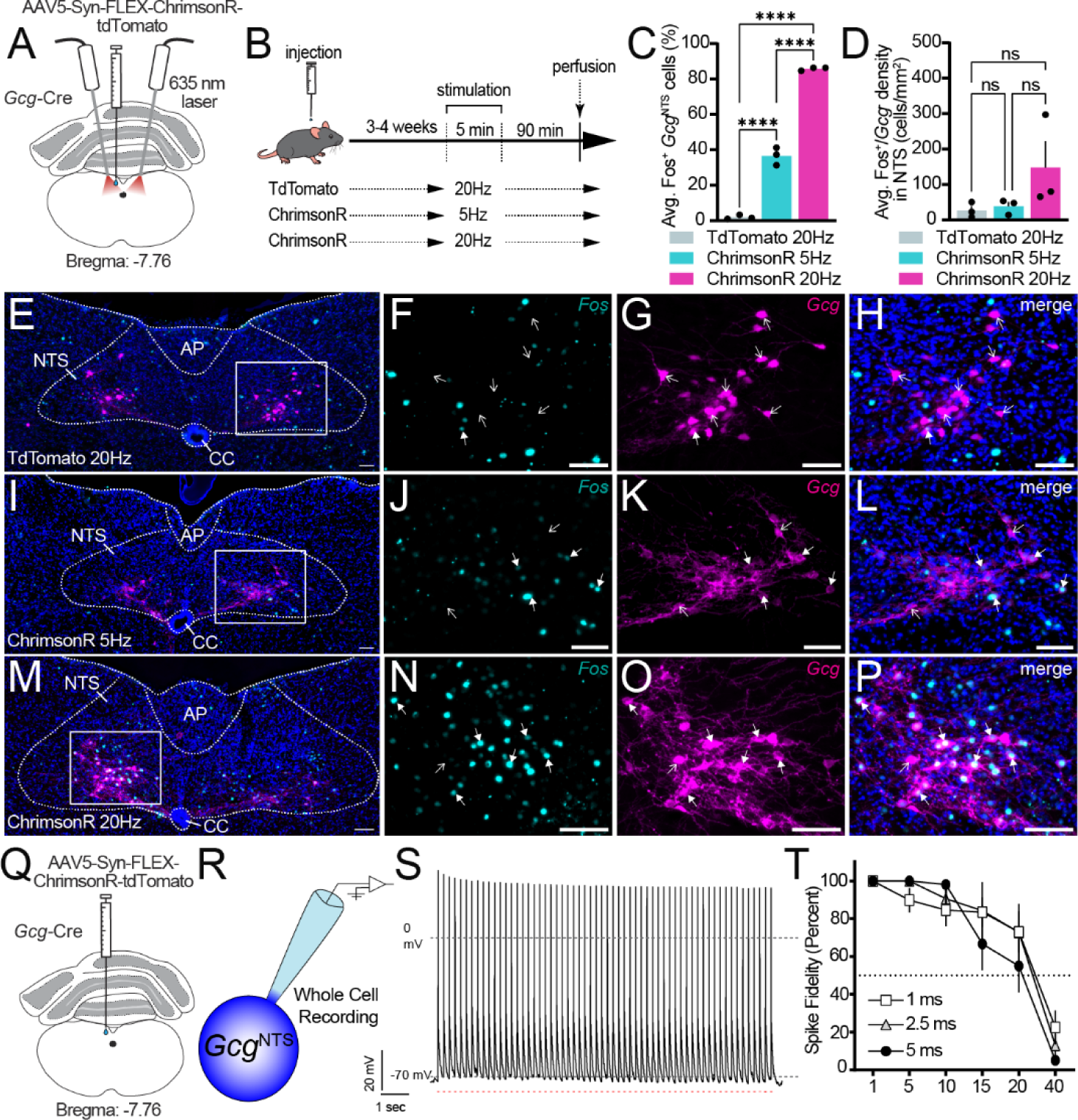
Tunable activation of *Gcg*^NTS^ neurons *in vivo* and *ex vivo* using ChrimsonR. (**A**) Surgical schematic of virus injection, optical fiber implant location, and 635 nm laser stimulation. (**B**) Schematic of experimental timeline and groups. (**C**) Average proportion of *Gcg*^NTS^ neurons expressing *Fos* after optical stimulation using male *Gcg*-iCre mice (n = 12; 2-4 images/mouse). Data analyzed using an ordinary one-way ANOVA followed by a Tukey’s multiple comparisons test. F (2, 6) = 579.2, *p* < 0.0001, for all comparisons: *p* < 0.0001. (**D**) Average density (cells/mm^2^) *Fos* expression in non-*Gcg*^NTS^ neurons. Data analyzed using an ordinary one-way ANOVA followed by a Tukey’s multiple comparisons test. F (2, 6) = 2.244, *p* = 0.1872, TdTomato 20Hz vs ChrimsonR 5 Hz: *p* = 0.9822, TdTomato 20 Hz vs. ChrimsonR 20 Hz: *p* = 0.2138, ChrimsonR 5 Hz vs ChrimsonR 20 Hz: *p* = 0.2675. (**E**) Large representative image of a TdTomato expressing NTS tissue section exposed to 20 Hz stimulation. White box illustrates close up view shown in (**F-H**) where (**F**) shows *Fos*+ cells (cyan), (**G**) shows *Gcg*+ cells (magenta), and (**H**) shows a merged view. (**I**) Large representative image of a ChrimsonR expressing NTS tissue section exposed to 5 Hz stimulation. White box illustrates magnified view shown in (**J-L**) where (**J**) shows *Fos*+ cells (cyan), (**K**) shows *Gcg*+ cells (magenta), and (**L**) shows a merged view. (**M**) Large representative image of a ChrimsonR expressing NTS tissue section exposed to 20 Hz stimulation. White box illustrates magnified view shown in (**N-P**) where (**N**) shows *Fos*+ cells (cyan), (**O**) shows *Gcg*+ cells (magenta), and (**P**) shows a merged view. For **E-P**, scale bar = 100µm. For **F-H**, **J-L**, and **N-P**, closed arrows indicate *Gcg*+/*Fos*+ cells and open arrows indicate *Gcg*+/*Fos*-cells. (**Q**) Schematic of stereotaxic surgery for electrophysiological experiments. (**R**) Whole cell patch clamp electrophysiological recording schematic for panels **S** and **T**. (**S**) Example trace of *Gcg*^NTS^ neuron expressing ChrimsonR held in current clamp at −70 (hyperpolarizing current injected) to reduce spontaneous activity at baseline. A train of 60 624 nm pulses (5 ms in length) was delivered through the microscope objective producing reliable action potentials. One sweep is shown. (**T**) Assessment of spike fidelity of *Gcg*^NTS^ neurons in response to 624 nm light pulses across increasing frequencies and pulsewidths of 1, 2.5, and 5 ms. Data were analyzed using two-way repeated measures ANOVA. Pulsewidth X Frequency: F (10, 125) = 1.392, *p =* 0.1892; Frequency: F (2.088, 52.20) = 58.75, *p* < 0.0001; Pulsewidth: F (2, 25) = 0.3035, *p* = 0.7409.

To demonstrate the impact of ChrimsonR on the neurophysiology of *Gcg*^NTS^ neurons, we used whole cell patch clamp electrophysiology in *ex vivo* NTS slices. We patched ChrimsonR-expressing *Gcg*^NTS^ neurons in NTS slices, providing trains of red light illumination while varying pulsewidth length and frequency to assess the spike fidelity of *Gcg*^NTS^ neurons in response to red light (**FIG. 4Q-S**). We observed some loss of spike fidelity after 10 Hz stimulation that was improved by using shorter pulsewidths that did not reach statistical significance. On average, we still observed greater than 50% spike fidelity at frequencies up to 20 Hz but observed a dramatic loss of spike fidelity at 40 Hz (**FIG. 4T**) irrespective of the pulsewidth. Overall, however, our data demonstrates that ChrimsonR is an effective optogenetic actuator for reversibly activating *Gcg*^NTS^ neurons *ex vivo* and *in vivo*.

### Optogenetic activation of Gcg^NTS^ neurons produces sex-specific anxiety-like behavior

Food intake is potentially altered by changes in overall locomotion and affective states, so we measured the behavioral consequences of optogenetic activation of *Gcg*^NTS^ neurons on locomotion and anxiety-like behavior using an open-field test where red light illumination was delivered in a reversible 3 min ON-OFF pattern at 20 Hz. We observed a significant difference in the pattern of locomotion, but not distance traveled, with all sexes (**FIG. 5A, 5D**). Interestingly, in a separate analysis of locomotion comparing OFF versus ON epochs, laser illumination seemed to subtly increase locomotion in both TdTomato and ChrimsonR groups (**FIG. 5E**). Separate analyses of optogenetic effects on locomotion revealed a similar overall effect in males, where there was no significant difference between groups in the overall distance traveled but a subtle effect of laser illumination on locomotion (**FIG. 5B, F, G**). We observed no differences in the pattern of distance traveled, overall, or as a function of laser epoch in female mice (**FIG. 5C, H, I**). These data demonstrate that optogenetic activation of *Gcg*^NTS^ neurons has no specific effect on locomotion. The open field assay has utility to gauge anxiety-like behavior, capitalizing on natural mouse thigmotaxis behaviors. Thus, we assessed center time as a function of transgene group and laser. For all mice and males, we observed no significant interaction between laser stimulation and ChrimsonR/TdTomato group (**FIG. 5J-K, M-P**). However, analysis of female mice revealed an overall reduction in center time that persisted during laser OFF epochs (**FIG. 5L, Q, R**). These data demonstrate that optogenetic stimulation of *Gcg*^NTS^ neurons is sufficient to produce sex-specific anxiety-like behavior that persists beyond laser stimulation epochs.

**Figure 5.**
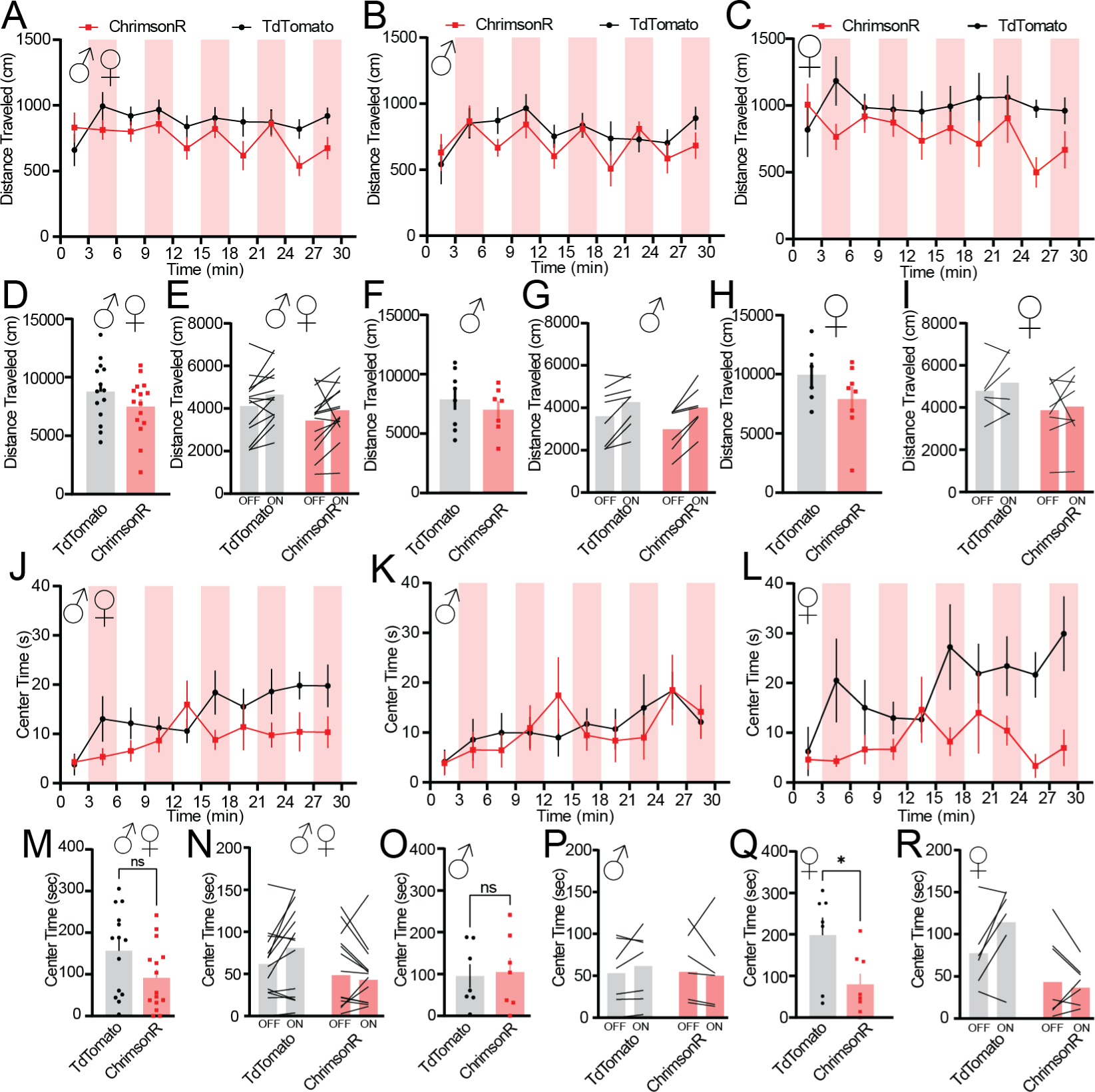
Open field behavior during optogenetic activation of *Gcg*^NTS^ neurons (**A**) Distance travelled in all sexes in the open field arena. Group (ChrimsonR/TdTomato) X Laser/Time Interaction effect: F (9, 234) = 2.112, *p* = 0.0294; Group effect: F (1, 26) = 3.052, *p* = 0.0924; Laser effect: F (2.717, 70.64) = 3.136, *p* = 0.0351. No significant post-tests. (**B**) Distance traveled for males only. Group (ChrimsonR/TdTomato) X Laser/Time Interaction effect: F (9, 108) = 1.403, *p* = 0.1959; Group effect: F (1, 12) = 1.494, *p* = 0.2450; Laser effect: F (2.220, 26.64) = 4.285, *p* = 0.0212. (**C**) Distance traveled for females only. Group (ChrimsonR/TdTomato) X Laser/Time Interaction effect: F (9, 108) = 1.403, *p* = 0.1959; Group effect: F (9, 108) = 1.656, *p =* 0.1937; Laser effect: F (2.639, 31.66) = 1.036, *p* = 0.3831. (**D**) Overall distance traveled in all sexes, *p* = 0.1865. (**E**) Epoch of distance traveled for all sexes. Group (ChrimsonR/TdTomato) X Laser/Time Interaction effect: F (9, 108) = 1.403, *p* = 0.8179; Group effect: F (1, 25) = 3.390, *p =* 0.0775; Laser effect: F (1, 25) = 8.869, *p* = 0.0064. (**F**) Overall distance traveled in males, *p* = 0.2450. (**G**) Epoch of distance traveled in males. Group (ChrimsonR/TdTomato) X Laser/Time Interaction effect: F (1, 12) = 0.9491, *p* = 0.3492; Group effect: F (1, 12) = 1.494, *p =* 0.2450; Laser effect: F (1, 12) = 30.65, *p* = 0.0001. (**H**) Overall distance traveled in females, *p* = 0.1937. (**I**) Epoch of distance traveled in females. Group (ChrimsonR/TdTomato) X Laser/Time Interaction effect: F (1, 12) = 0.1229, *p* = 0.7319; Group effect: F (1, 12) = 1.896, *p =* 0.1937; Laser effect: F (1, 12) = 0.8368, *p* = 0.3783. (**J**) Center time across all sexes. Group (ChrimsonR/TdTomato) X Laser/Time Interaction effect: F (9, 234) = 1.711, *p* = 0.0873; Group effect: F (1, 26) = 3.468, *p =* 0.0739; Laser effect: F (3.369, 87.59) = 3.409, *p* = 0.0171. No significant post-tests. (**K**) Center time for males only. Group (ChrimsonR/TdTomato) X Laser/Time Interaction effect: F (9, 110) = 0.6419, *p* = 0.7590; Group effect: F (1, 14) = 0.1541, *p =* 0.7006; Laser effect: F (2.737, 33.45) = 2.619, *p* = 0.0718. (**L**) Center time for females only. Group (ChrimsonR/TdTomato) X Laser/Time Interaction effect: F (9, 108) = 1.863, *p* = 0.0652; Group effect: F (1, 12) = 6.871, *p =* 0.0223; Laser effect: F (3.211, 38.53) = 2.063, *p* = 0.1173. (**M**) Overall center time in all sexes, *p* = 0.0631. (**N**) Center time by epoch in all sexes. Group (ChrimsonR/TdTomato) X Laser/Time Interaction effect: F (1, 27) = 5.278, *p* = 0.0296; Group effect: F (1, 27) = 2.710, *p =* 0.1113; Laser effect: F (3.211, 38.53) = 2.063, *p* = 0.2240. (**O**) Overall center time in males, p = 0.8322. (**P**) Center time by epoch in males. Group (ChrimsonR/TdTomato) X Laser/Time Interaction effect: F (1, 12) = 1.158, *p* = 0.3031; Group effect: F (1, 12) = 0.04877, *p =* 0.8289; Laser effect: F (1, 12) = 0.09107, *p* = 0.7680. (**Q**) Overall center time in females, *p* = 0.0271. (**R**) Epoch of center time in females. Group (ChrimsonR/TdTomato) X Laser/Time Interaction effect: F (1, 12) = 5.738, *p* = 0.0338; Group effect: F (1, 12) = 6.871, *p =* 0.0223; Laser effect: F (1, 12) = 2.770, *p* = 0.1219. For all panels, n = 15 ChrimsonR mice (7 male, 8 female), n = 13 TdTomato mice (7 male, 6 female). For D, F, H, M, O, and Q, data were analyzed using unpaired student *t* tests. For A-C, E, G, I, J-L, N, P, and R data were analyzed using two-way Repeated Measures ANOVA with Sidak’s multiple comparisons of TdTomato vs ChrimsonR. Red laser epochs represented by the red fill bar.

### Optogenetic stimulation of Gcg^NTS^ neurons is sufficient to produce real-time place avoidance

Activation of GLP-1Rs is known to exert broad effects on motivation and consumption of natural and drug rewards.^68^ To assess the impact of *Gcg*^NTS^ neuron stimulation on motivation and valence, we combined optogenetic stimulation of *Gcg*^NTS^ neurons with real time place conditioning. In this assay, mice explore a chamber with two identical sides where only one side is paired with laser stimulation. At frequencies of 1, 5, and 10 Hz, we observed no significant preference or aversion to the laser-paired side (**FIG. 6A-C, G**). However, high frequency optogenetic stimulation of *Gcg*^NTS^ neurons was sufficient to produce aversion and avoidance behavior in both sexes of mice (**FIG. 6A-C**). We observed that 20 Hz stimulation was sufficient to produce aversion in female mice but not male mice, but a direct comparison of male and female ChrimsonR mice at 20 Hz stimulation did not reach statistically significance (*p* = 0.0820). We also observed a significant difference in the distance traveled between TdTomato and ChrimsonR mice at high frequencies of 20 and 40 Hz (**FIG. 6D-F**), which is consistent with a reduction in overall exploration after learning avoidance behavior (**FIG. 6G).** To confirm that activation of *Gcg*^NTS^ neurons is sufficient to produce real time learning and that this is not a learned side preference, we ran a separate session at 40 Hz that included a reversal of the laser-paired side at the midpoint of the assay. ChrimsonR mice displayed a significant aversion to the laser-paired side even after reversal (**FIG. 6H**). Together, these data show that high frequency activation of *Gcg*^NTS^ neurons is sufficient to serve as a teaching signal and produce avoidance behavior.

**Figure 6.**
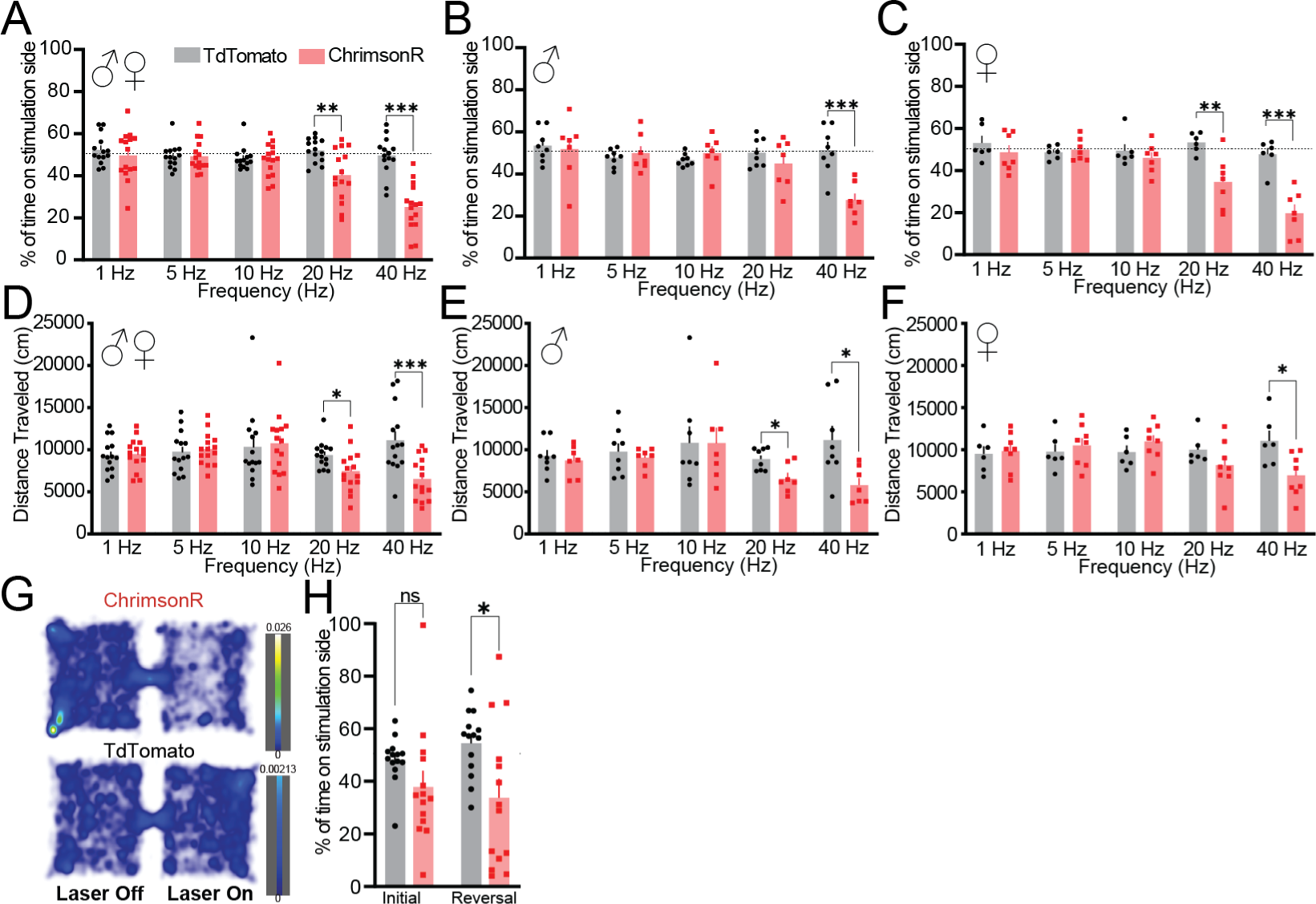
Activation of *Gcg*^NTS^ neurons produces real-time place avoidance. (**A**) Percent of time spent on the stimulation side for all sexes across frequencies (1, 5, 10, 20, and 40 Hz). Unpaired Student’s *t* tests: 1 Hz, *p* = 0.4362; 5 Hz, *p* = 0.9972; 10 Hz, *p* = 0.7499; 20 Hz, *p* = 0.0031; 40 Hz. *p* < 0.0001. (**B**) Percent of time spent on the stimulation side for male mice only across all frequencies. Unpaired Student’s *t* tests: 1 Hz, *p* = 0.7801; 5 Hz, *p* = 0.5364; 10 Hz, *p* = 0.3056; 20 Hz, *p* = 0.3029; 40 Hz, *p* = 0.0005. (**C**) Percent of time spent on the stimulation side for female mice only across frequencies. Unpaired Student’s *t* tests: 1 Hz, *p* = 0.3914; 5 Hz, *p* = 0.6348; 10 Hz, *p* = 0.4202; 20 Hz, *p* = 0.0050; 40 Hz, *p* = 0.0002. (**D**) Distance traveled during real time place conditioning for all sexes across frequencies. Unpaired Student’s *t* tests: 1 Hz, *p* = 0.8618; 5 Hz, *p* = 0.7122; 10 Hz, *p* = 0.7844; 20 Hz, *p* = 0.0210; 40 Hz, *p* = 0.0008. (**E**) Distance traveled during real time place conditioning for male mice only across frequencies. Unpaired Student’s *t* tests: 1 Hz, *p* = 0.5798; 5 Hz, *p* = 0.5450; 10 Hz, *p* = 0.9911; 20 Hz, *p* = 0.0100; 40 Hz, *p* = 0.0168. (**F**) Distance traveled during real time place conditioning for female mice only across frequencies. Unpaired Student’s *t* tests: 1 Hz, *p* = 0.7830; 5 Hz, *p* = 0.5625; 10 Hz, *p* = 0.2658; 20 Hz, *p* = 0.2036; 40 Hz, *p* = 0.0162. (**G**) Heat map revealing ChrimsonR animals localizing on the non-laser paired side of the arena, while TdTomato injected mice constantly moved across both sides of the arena. (**H**) Percent time spent in stimulation side using a reversal paradigm at 40 Hz for all sexes. F (1, 26) = 0.8631, *p* = 0.3614; Initial TdTomato vs. ChrimsonR: *p* = 0.2963, Reversal TdTomato vs. ChrimsonR: *p* = 0.0120. For all panels, n = 15 ChrimsonR mice (7 male, 8 female), n = 13 TdTomato mice (7 male, 6 female). For **A**-**F** data were analyzed using unpaired Student’s *t* tests for TdTomato vs. ChrimsonR. For **H** data were analyzed using two-way repeated measures ANOVA with Sidak’s multiple comparisons test.

### Reversible optogenetic activation of Gcg^NTS^ neurons reduces feeding and appetitive behavior

Several published studies have demonstrated that activation of *Gcg^NTS^* neurons reduces food intake using chemogenetics ^31, 38, 39, 52^, but these studies lack the ability to tune stimulation, reversibly alter *Gcg*^NTS^ neuron stimulation, or to pair stimulation with certain behavioral epochs (a la closed loop). As a first effort, we tested whether optogenetic activation is sufficient to reduce food intake using a refeed design similar to our electrophysiological paradigm. Mice were trained to use <u>F</u>eeding <u>E</u>xperimental <u>D</u>evices for these studies so that we could clearly map pellet retrievals to optogenetic stimulation in a granular fashion (**FIG. 7A-B**).^60, 61^ After training in the absence of stimulation, mice were food deprived overnight. The next day refeeding was conducted where optogenetic stimulation was provided in a reversible fashion (5 min ON-OFF) counterbalanced across animals and groups. In all mice, we observed no significant difference in pellets retrieved during OFF vs ON across 1, 5, and 10 Hz stimulation of *Gcg*^NTS^ neurons (**FIG. 7C-E**), but reversible 20 Hz stimulation produced a significant difference in pellets retrieved (**FIG. 7F**) relative to TdTomato controls which surprisingly showed a significant increase. When analyzed separately, reversible optogenetic stimulation of *Gcg*^NTS^ neurons reduced pellet retrievals at 5 and 20 Hz in male mice (**FIG. 7G-J**). Female mice did not display a specific and significant reduction in pellet retrievals at any frequency with the 20 Hz session not meeting our threshold for statistical significance (**FIG. 7K-N**). Nonetheless an anorexigenic trend was evident. These data demonstrate that, like chemogenetic actuators, reversible optogenetic activation of *Gcg*^NTS^ neurons using ChrimsonR can significantly reduce food intake.

**Figure 7.**
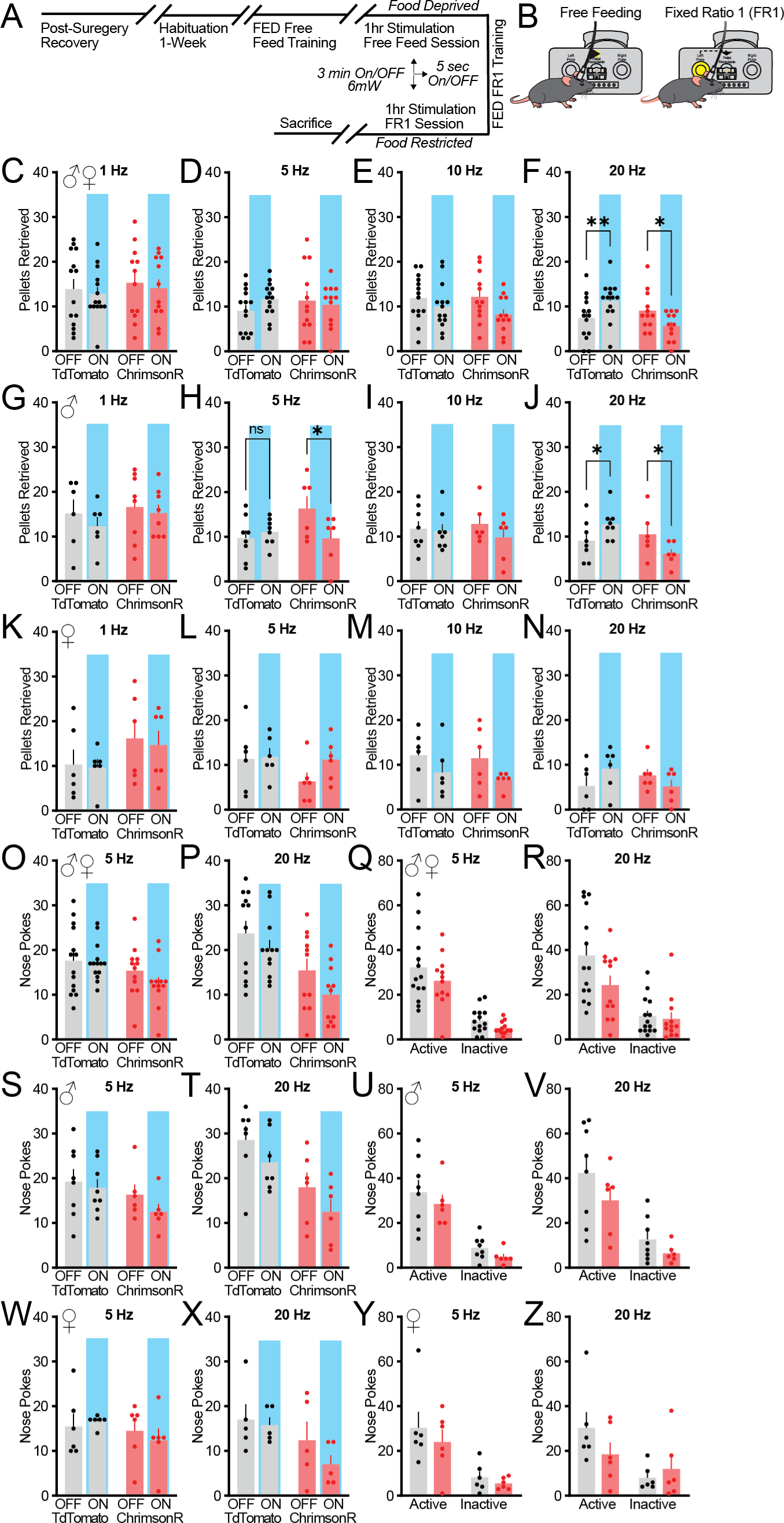
Impact of optogenetic activation of *Gcg*^NTS^ neurons on feeding and appetitive behavior. (**A**) Timeline sequence of Refeed and Fixed Ratio 1 (FR1) experiments, showing a reversible paradigm. (**B**) Illustration of (left) free feeding and (right) FR1 paradigms using an experimental feeding device (FED). All experiments included ChrimsonR (n = 13-14; 6 male, 7-8 female) and TdTomato (n = 10-13; 6-7 male, 4-6 female) mice. For **C-N**, the number of pellets retrieved during the OFF/ON epochs of laser stimulation at various frequencies (ON epoch highlighted with blue background on bar charts). (**C**) Pellets retrieved at 1 Hz for all mice. Group (ChrimsonR/TdTomato) X Laser/Epoch effect: F (1, 24) = 0.00423, *p* = 0.9487; Group effect: F (1, 24) = 0.2695, *p =* 0.6084; Laser/Epoch effect: F (1, 24) = 0.7148, *p =* 0.4062. (**D**) Pellets retrieved at 5 Hz for all mice. Group (ChrimsonR/TdTomato) X Laser/Epoch effect: F (1, 24) = 1.554, *p* = 0.2245; Group effect: F (1, 24) = 0.09446, *p =* 0.7612; Laser effect: F (1, 24) = 0.3655, *p =* 0.5511. (**E**) Pellets retrieved at 10 Hz for all mice. Group (ChrimsonR/TdTomato) X Laser/Epoch effect: F (1, 24) = 0.7721, *p* = 0.3883; Group effect: F (1, 24) = 0.2205, *p =* 0.6429; Laser effect: F (1, 24) = 6.674, *p =* 0.0163. (**F**) Pellets retrieved at 20 Hz for all mice. Group (ChrimsonR/TdTomato) X Laser/Epoch effect: F (1, 24) = 20.56, ***p* = 0.0001**; Group effect: F (1, 24) = 1.835, *p =* 0.1882; Laser effect: F (1, 24) = 0.05398, *p =* 8182; Tdtomato OFF vs ON, ***p* = 0.0036**; ChrimsonR OFF vs ON, ***p* = 0.0146**. (**G**) Pellets retrieved at 1 Hz for male mice. Group (ChrimsonR/TdTomato) X Laser/Epoch effect: F (1, 12) = 0.1166, *p* = 0.7387; Group effect: F (1, 12) = 0.5806, *p =* 0.4608; Laser effect: F (1, 12) = 0.9706, *p =* 0.3440. (**H**) Pellets retrieved at 5 Hz for male mice. Group (ChrimsonR/TdTomato) X Laser/Epoch effect: F (1, 12) = 6.534, ***p* = 0.0252**; Group effect: F (1, 12) = 1.313, *p =* 0.2741; Laser effect: F (1, 12) = 0.2829, *p =* 0.1184. TdTomato OFF vs. ON: *p* = 0.7667, ChrimsonR OFF vs. ON: ***p* = 0.0316**. (**I**) Pellets retrieved at 10 Hz for male mice. Group (ChrimsonR/TdTomato) X Laser/Epoch effect: F (1, 12) = 0.6875, *p* = 0.4232; Group effect: F (1, 12) = 0.00669, *p =* 0.9362; Laser effect: F (1, 12) = 1.348, *p =* 0.2683. (**J**) Pellets retrieved at 20 HZ for male mice. Group (ChrimsonR/TdTomato) X Laser/Epoch effect: F (1, 12) = 19.94; ***p* = 0.0008**. Group effect: F (1, 12) = 1.825, *p =* 0.2017; Laser effect: F (1, 12) = 0.1038, *p =* 0.7528. TdTomato OFF vs. ON: ***p* = 0.0162**, ChrimsonR OFF vs. ON: ***p* = 0.0162**. (**K**) Pellets retrieved at 1 Hz for female mice. Group (ChrimsonR/TdTomato) X Laser/Epoch effect: F (1, 10) = 0.04463; *p* = 0.8369. Group effect: F (1, 10) = 0.1.765, *p =* 0.2135; Laser effect: F (1, 10) = 0.3017, *p =* 0.5949. (**L**) Pellets retrieved at 5 Hz for female mice. Group (ChrimsonR/TdTomato) X Laser/Epoch effect: F (1, 10) = 0.9230; *p* = 0.3593. Group effect: F (1, 10) = 1.621, *p =* 0.2318; Laser effect: F (1, 10) = 1.398, *p =* 0.2644. (**M**) Pellets retrieved at 10 Hz for female mice. Group (ChrimsonR/TdTomato) X Laser/Epoch effect: F (1, 10) = 0.08072; *p* = 0.7821. Group effect: F (1, 10) = 0.1939, *p =* 0.6691; Laser effect: F (1, 10) = 6.063, ***p =* 0.0335**. (**N**) Pellets retrieved at 20 Hz for female mice. Group (ChrimsonR/TdTomato) X Laser/Epoch effect: F (1, 10) = 4.885; *p* = 0.0515. Group effect: F (1, 10) = 0.1672, *p =* 0.6912; Laser effect: F (1, 10) = 0.2165, *p =* 0.6517. (**O**) The number of nose pokes during OFF/ON epochs at 5 Hz for all mice. Group (ChrimsonR/TdTomato) X Laser/Epoch effect: F (1, 24) = 1.563; *p* = 0.2232. Group effect: F (1, 24) = 3.071, *p =* 0.0925; Laser effect: F (1, 24) = 2.291, *p =* 0.1432. (**P**) The number of nose pokes during OFF/ON epochs at 20 Hz for all mice. Group (ChrimsonR/TdTomato) X Laser/Epoch effect: F (1, 21) = 0.8292; *p* = 0.3728. Group effect: F (1, 21) = 8.743, ***p =* 0.0075**; Laser effect: F (1, 21) = 15.71, ***p =* 0.0007**. (**Q**) The number of active and inactive nose pokes at 5 Hz for all mice. Group (ChrimsonR/TdTomato) X Side Choice effect: F (1, 24) = 0.2343, *p* = 0.6328; Group effect: F (1, 24) = 2.306, *p =* 0.1420; Side Choice effect: F (1, 24) = 67.92, ***p* < 0.0001**. (**R**) The number of active and inactive nose pokes at 20 Hz for all mice. Group (ChrimsonR/TdTomato) X Side Choice effect: F (1, 24) = 2.213, *p* = 0.1499; Group effect: F (1, 24) = 3.529, *p =* 0.0725; Side Choice effect: F (1, 24) = 27.36, ***p* < 0.0001**. (**S**) The number of nose pokes during OFF/ON epochs at 5 Hz for male mice. Group (ChrimsonR/TdTomato) X Laser/Epoch effect: F (1, 12) = 0.7341; *p* = 0.4083. Group effect: F (1, 12) = 1.906, *p =* 0.1925; Laser effect: F (1, 12) = 3.295, *p =* 0.0945. (**T**) The number of nose pokes during OFF/ON epochs at 20 Hz for male mice. Group (ChrimsonR/TdTomato) X Laser/Epoch effect: F (1, 11) = 0.02696; *p* = 0.8725. Group effect: F (1, 11) = 7.866, ***p =* 0.0171**; Laser effect: F (1, 11) = 11.89, ***p =* 0.0054**. (**U**) The number of active and inactive nose pokes at 5 Hz for male mice. Group (ChrimsonR/TdTomato) X Side Choice effect: F (1, 12) = 0.03337, *p* = 0.8581; Group effect: F (1, 12) = 1.159, *p =* 0.3029; Side Choice effect: F (1, 12) = 53.85, ***p* < 0.0001**. (**V**) The number of active and inactive nose pokes at 20 Hz for male mice. Group (ChrimsonR/TdTomato) X Side Choice effect: F (1, 12) = 0.2772, *p* = 0.6081; Group effect: F (1, 12) = 2.759, *p =* 0.1226; Side Choice effect: F (1, 12) = 21.37, ***p* = 0.0006**. (**W**) The number of nose pokes during OFF/ON epochs at 5 Hz for female mice. Group (ChrimsonR/TdTomato) X Laser/Epoch effect: F (1, 10) = 0.9217; *p* = 0.3597. Group effect: F (1, 10) = 0.8418, *p =* 0.3805; Laser effect: F (1, 10) = 0.08295, *p =* 0.7792. (**X**) The number of nose pokes during OFF/ON epochs at 20 Hz for female mice. Group (ChrimsonR/TdTomato) X Laser/Epoch effect: F (1, 8) = 1.521; *p* = 0.2525. Group effect: F (1, 8) = 2.896, *p =* 0.1272; Laser effect: F (1, 8) = 3.755, *p =* 0.0886. (**Y**) The number of active and inactive nose pokes at 5 Hz for female mice. Group (ChrimsonR/TdTomato) X Side Choice effect: F (1, 10) = 0.1507, *p* = 0.7060; Group effect: F (1, 10) = 0.8596, *p =* 0.3757; Side Choice effect: F (1, 10) = 18.54 ***p* = 0.0015**. (**Z**) The number of active and inactive nose pokes at 20 Hz for female mice. Group (ChrimsonR/TdTomato) X Side Choice effect: F (1, 10) = 2.125, *p* = 0.1756; Group effect: F (1, 10) = 0.5298, *p =* 0.4834; Side Choice effect: F (1, 10) = 7.048, *p* = **0.0241**. For **C-Z**, data were analyzed using two-way repeated measures ANOVA with Sidak’s multiple comparisons.

Previous studies have not addressed how the activation of *Gcg*^NTS^ neurons impacts food-seeking appetitive behavior. Under food restriction, we trained optogenetic cohorts to nose poke to receive a chocolate sucrose pellet using a fixed ratio 1 design using FEDs (**FIG. 7A**). After training, we focused experimental sessions on one low (5 Hz) and one high (20 Hz) activation frequency where stimulation was provided in counterbalanced 5 min OFF/ON epochs as in the refeeding study. At 5 Hz, we did not observe a significant reduction in overall active port nose pokes as a function of group or laser epoch, although active port nose pokes were reduced (**FIG. 7O, Q**). In this session, mice in both groups showed a clear preference, showing instrumental responses for the active nose poke (**FIG. 7Q**). When analyzed separately by sex, we observed similar overall trends as the grouped data but overall there was no significant reduction in appetitive nose pokes overall or as a function of laser epoch (**FIG. 7S, U, W, Y**). For the 20 Hz session, again we observed a clear preference for the active port relative to the inactive port (**FIG. 7R**), and, similar to 5 Hz, there was an overall reduction in poking between groups that did not reach significance. Separate analyses by sex reveal similar overall trends with both male and female mice showing a preference for the active port but no significant differences overall (**FIG. 7V, Z**). As a function of laser epoch, we observed no significant interaction between transgene group and the laser epoch. However, we observed a significant main effect of group demonstrating that optogenetic stimulation of *Gcg*^NTS^ neurons produces reductions in active port nose poking that significantly differ between ChrimsonR and TdTomato controls but persist outside of laser stimulation epochs (**FIG. 7P**). Separate analyses by sex revealed this same phenomenon in males, but female mice were not significantly different as a function of group but displaying the same trend (**FIG. 7T,X**).

## DISCUSSION

In this study, we made some important advancements in our understanding of the brain’s GLP-1-producing neurons, *Gcg*^NTS^ neurons.

Previous studies using rats and mice identify the presence of GLP-1 and *Gcg* in the NTS, however a rigorous comparison of the density and distribution of *Gcg*^NTS^ neurons in male and female animals had not been performed.^63,69,43,70,71,42,41,38,44^ Here we show in mice that there is no difference in the density or distribution of *Gcg*^NTS^ neurons between male and female mice. We also provide validation at an mRNA level of a published *Gcg-*iCre driver transgenic mouse and that this mouse provides robust and faithful access to *Gcg*^NTS^ neurons, though our data show that it’s possible that we are not capturing the complete *Gcg*^NTS^ neuron population.^56^ Previous studies had also demonstrated that *Gcg*^NTS^ neurons are anatomically distinct from neighboring noradrenergic neurons^49, 63^, which we independently validate in this study.

We adapted a published paradigm in rats combined with *Fos* immunohistochemistry to examine the recruitment of the *Gcg*^NTS^ neuron system to eating.^42^ Overall we did not observe a significant activation of *Gcg*^NTS^ neuron in response to multiple levels of food intake. However, we were limited in our statistical power and may not have a large enough sample to make definitive conclusions. When we examined the relationship between food intake and Fos activation of *Gcg*^NTS^ neurons we observed a clear linear relationship as has been previously noted.^41, 42^ In comparing our study with previous ones, we speculate that the mice in our AL sample may not have reached the levels of intake needed to demonstrate Fos activation. Also, the previous studies used a liquid diet which may provide more enhance gastric interoceptive input to the NTS. The differences in these paradigms are worth exploring in a larger sample of animals.

Only two studies to date have examine the neurophysiology of *Gcg*^NTS^ neurons.^49, 50^ Despite some technical differences in our recording mode (perforated patch vs whole/cell attached) and the use of a different transgenic mouse line, we made very similar observations overall. First, under *ab libitum* conditions all *Gcg*^NTS^ neurons display spontaneous action potentials as we observed via the cell-attached method where, on average, *Gcg*^NTS^ neurons fire at 2-3 Hz but can range from >0 – 7 Hz. This property was not significantly different between male and female mice. Hisadome *et* al. observed that *Gcg*^NTS^ neurons fire at 1.5 or 1.9 Hz via perforated patch but also observed some silent *Gcg*^NTS^ neurons. In our hands, we observed that spontaneous action potentials in *Gcg*^NTS^ neurons was very prone to rundown (not shown) using whole-cell recordings. Action potential rundown has been observed in another NTS population.^72^ We speculate that even perforation may lead to subtle rundown of *Gcg^NTS^* neuron firing, which is why we used cell-attached. Interestingly, we observed that 24-hour food deprivation reduces spontaneous firing of *Gcg*^NTS^ neurons and this depended on the sex of the animal, while refeeding restored the mean firing rate to AL levels but did not increase it. An independent assessment of the distribution, however, revealed that refeeding produces a significant difference in the overall distribution of *Gcg*^NTS^ neuron firing rates. We observed individual neurons from refed mice firing at ∼9 Hz. The *ex vivo* slice prep gives us new and important insight but is limited in its ability to capture transient changes in activity. Future studies using *in vivo* approaches will be necessary to gain a better understanding of transient changes in activity that occur prior to, during, and after food intake.

We also observed a very pronounced shift in the excitability of *Gcg*^NTS^ neurons using whole cell recording methodology. Food deprivation led to a significant decrease in membrane resistance, suggesting greater ionic conductance or more open ion channels. The source ion channels are not known at this time, but we speculate that there may be differences in the magnitude of voltage-gated potassium channel conductance as a function of energy status.^73^ Food deprivation also resulted in a significant increase in the rheobase and a rightward shift in the magnitude of action potential firing in response to depolarizing currents. In a separate analysis, we also observed that there are sex differences in the sensitivity of depolarizing currents to elicit action potentials, where *Gcg*^NTS^ neurons from female mice are less sensitive and have reduced action potential throughput (max firing rate). Overall these data showing decreased *Gcg*^NTS^ neuron firing and excitability are consistent with previous studies in rats showing that food deprivation can decrease the overall *Fos* levels of GLP-1+ NTS neurons.^70, 71^ Future studies will investigate the molecular mechanisms of these changes in physiology, but’s also possible that there are changes in excitation/inhibition as a function of energy state that accompany these intrinsic differences.^74^

To better understand the behavioral contribution of *Gcg*^NTS^ neurons, we pioneered the use of optogenetics to manipulate these neurons *ex vivo* and *in vivo*. To set the stage for future experiments combining optogenetic manipulation with GCaMP recordings, we used the red light-gated opsin ChrimsonR.^67^ We found that ChrimsonR was very effective at evoking action potentials using an *ex vivo* slice preparation, although we observed a dramatic loss of spike fidelity at frequencies greater than 20 Hz. This phenomenon may be due to the T_off_ rate of the opsin, but other studies using ChrimsonR have observed higher spike fidelities at greater frequencies relative to ones we used. Its more likely that there are inherent limitations to the peak spike rate of *Gcg*^NTS^ neurons due to its ion channel composition. Interestingly, our excitability data may also support this idea, but it requires further testing. We show here that red-light activation of ChrimsonR-expressing *Gcg^NTS^* neurons for 5 min can produce frequency-dependent, tunable levels of activation as readout by Fos. Twenty Hz stimulation produced a remarkable 85% activation of ChrimsonR-expressing *Gcg^NTS^* neurons. To perform these studies, we used a lower laser power than is typically used (6 mW).

Having established optogenetics as a tool to manipulate *Gcg*^NTS^ neurons, we investigated their causal behavioral role. Our first experiment established that optogenetic activation of *Gcg*^NTS^ neurons (at 20 Hz) produces no major changes in locomotion, although we noted a subtle effect of laser alone irrespective of group. In the open field, we also observed sex-dependent anxiety-like behavior where optogenetic stimulation of *Gcg*^NTS^ neurons reduced exploration time of the center but only in female mice. Its possible we were at a floor effect with our male control group to see this effect. However, in our next experiment we also observed that optogenetic activation of *Gcg*^NTS^ neurons produced a significant real-time place aversion. At 20 Hz, we observed aversion in our female, but not male mice. A direct head-to-head comparison of time spent in the laser-paired chamber between male and female mice did not reach significance, but displayed a nonsignificant trend that suggests sex differences in the ability of optogenetic stim of *Gcg*^NTS^ neurons to produce avoidance behavior. Overall, this is consistent with previous studies demonstrating that aversive stimuli like stress can activate *Gcg*^NTS^ neurons.^44, 69^ We look forward to future studies examining how optogenetic stimulation modulates passive and active avoidance to aversive stimuli and ethological stressors. Interestingly, the production of anxiety-like and avoidance behavior by optogenetic activation of *Gcg*^NTS^ neurons was achieved using reversible stimulation as opposed to constant activation. This implies that even transient, brief activation of *Gcg*^NTS^ neurons at high frequencies may have important behavioral consequences and that it can serve as a potent teaching signal. Although the endogenous firing rates of *Gcg*^NTS^ neurons are unknown, we speculate the firing rates of these neurons are higher *in vivo* with an intact glutamatergic vagus than they are *ex vivo* and that stimulation of these neurons up to 20 Hz is within a physiological range.

We used optogenetics to replicate previous studies demonstrating that activation of *Gcg*^NTS^ neurons reduces food intake.^31, 38, 39, 52^ Beyond these studies, we show that significant acute anorexigenic behavior as a consequence of *Gcg*^NTS^ neuron activation primarily occurs at high frequencies of 20 Hz. However, we do not exclude the possibility that our study was limited by potential floor effects that might reveal subtle decreases in intake that occur at lower frequencies of 5 and 10 Hz. Future studies using stimulation over longer assays with greater overall intake will be important to test this hypothesis. Importantly, our study demonstrates that transient and reversible activation of *Gcg*^NTS^ neurons can produce significant anorexigenic effects in mice. Our paradigm of 5 min OFF/ON along with alternating 5 sec of stimulation during ON periods (to prevent excessive heating) implies that activation of *Gcg*^NTS^ neurons during 25% of the potential assay was sufficient to reduce food intake. Future studies coupling stimulation of *Gcg*^NTS^ neurons in a closed-loop manner are needed to determine the relevant behavioral sequelae through which activation of *Gcg*^NTS^ neuron-mediated suppression of feeding operates.

Although several studies have demonstrated that GLP-1R agonists can reduce appetitive for natural and drug rewards^75–77^, the role of *Gcg*^NTS^ neurons in modulating appetitive behavior in addition to consumption is unknown. We combined optogenetic stimulation of *Gcg*^NTS^ neurons with operant food consumption using an FR1 task for chocolate sucrose pellets. At 5 Hz stimulation we did not observe a significant effect on appetitive nose poking for food either overall or as a function of the laser epoch, though a nonsignificant reduction in overall nose pokes was noted. Our limitations here are similar to our refeeding experiments. It is possible that a longer assay or persistent stimulation at 5 Hz might have revealed a significant effect. At 20 Hz we made some important and novel observation in that optogenetic activation of *Gcg*^NTS^ neurons produced significant reductions in active port nose pokes, but did not differ as a function of the laser being on. This is consistent with the idea that stimulation of *Gcg*^NTS^ neurons can produce lasting effects on overall appetitive behavior. Consistent with this idea, we also saw reductions in nose pokes to the inactive port but we are at a floor effect to see a significant reduction with optogenetic stim. Importantly, appetitive behavior at the end of training was identical between the two groups (data not shown). Further testing is needed under greater motivational demand to test this idea more completely.

Our study has important implications for our understanding the brain’s endogenous GLP-1 system and highlights how further study of this system is needed. Critically, we have a very limited view of how *Gcg*^NTS^ neurons interact with downstream GLP-1R expressing systems. These will be pivotal to developing better pharmacotherapies and understanding the ramifications of GLP-1R based therapeutics for the treatment of obesity and diabetes.

---

**Supplemental Figure 1.**
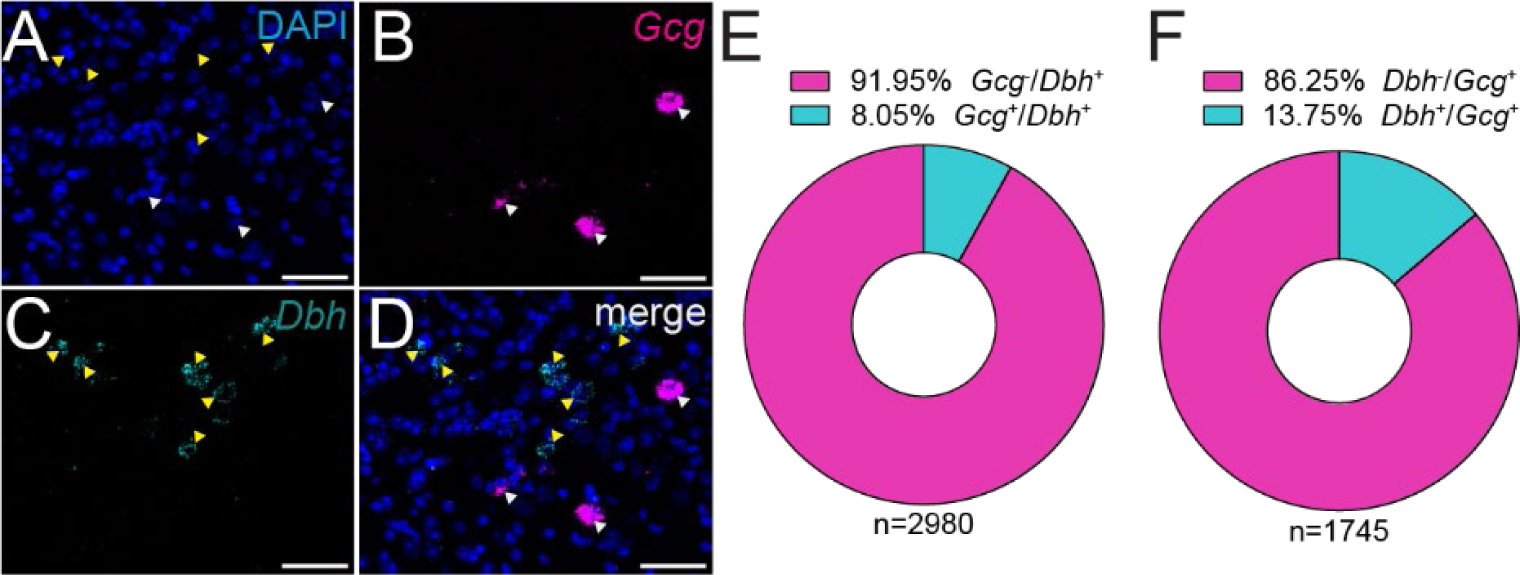
Minimal overlap of *Gcg* mRNA and *Dbh* mRNA expression in NTS neurons. (A-D) Single channel representative images showing enlarged view of (A) DAPI (blue), (B) *Gcg* mRNA expressing neurons (magenta), (C) *Dbh* mRNA expressing neurons (cyan), and (D) a merged view. Scale bar = 50µm. (E) Quantification of *Gcg* mRNA expression within *Dbh*+ cells (n = 2980 cells). (F) Quantification of *Dbh* mRNA expression within *Gcg*+ cells (n = 1745 cells). For E, the mean number of *Gcg*+ cells counted per image was 60.17 ± 4.01 (s.e.). For F, the mean number of *Dbh*+ cells counted per image was 102.8 ± 10.75 (s.e.). For E-F, data were obtained from 29 images taken from C57BL6/J mice (n = 3) with 8-11 images per mouse.

**Supplemental Figure 2.**
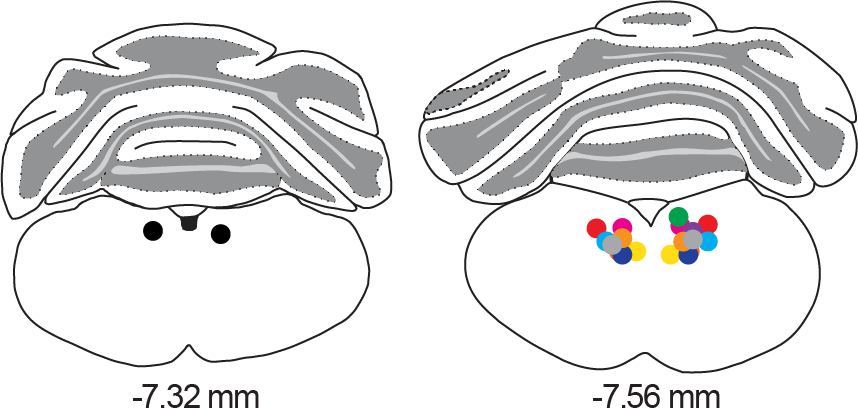
Histological validation of optical fiber placement for all mice used in optogenetic experiments. Optical fiber termination points were located between −7.32 and −7.56 mm with respect to bregma. Each dot color pair represents the optical fiber termination point of one mouse.

## Notes

### Competing Interest Statement

The authors have declared no competing interest.

